# Crystallographic Ensembles Reveal the Structural Basis of Binding Entropy in SARS-CoV2 Macrodomain

**DOI:** 10.1101/2025.11.25.690589

**Authors:** Louella Seo, Ian Farran, Ahmed Aslam, Xinyun Li, Priyadarshini Jaishankar, Alan Ashworth, James S. Fraser, Adam R. Renslo, Stephanie A. Wankowicz

## Abstract

Structure-based drug design has traditionally focused on optimizing static, enthalpic interactions between ligands and proteins or on displacing binding site solvent molecules to entropically favor binding. A potentially large contributor to binding thermodynamics is the difference in conformational entropy of the protein upon binding a ligand; however, this has been difficult to quantify especially in high throughput. Here, through multiconformer ensemble modeling of hundreds of ligand-bound SARS-CoV-2 Macrodomain (Mac1) X-ray structures, we show how ligand binding reorganizes both protein conformational entropy and water molecules. By applying an optimal transport–based clustering algorithm, we show how specific protein–ligand interactions patterns drive the magnitude and spatial redistribution of conformational entropy and solvent networks. Using isothermal titration calorimetry (ITC), we demonstrate a correlation between experimental binding thermodynamics and conformational entropy estimated from structural ensemble models, showing that increased conformational heterogeneity and a less connected hydrogen-bonded water network lead to more entropic binding. These results establish a framework for extracting thermodynamically meaningful information from crystallographic ensembles, enabling the integration of entropic effects into prospective, ensemble-aware drug discovery.

## Introduction

Structure-based drug design is a cornerstone of modern drug discovery. Currently, this approach relies largely on static interpretations of protein–ligand interactions, obscuring the reality that binding affinity emerges from both enthalpy, derived from specific interactions, and entropy, derived from the multiplicity of solvent, protein, and ligand conformational states^1,2.3,4^. The emphasis on enthalpic interactions is primarily due to the lack of quantitative or even qualitative measurements of entropy from the majority of experimental structural biology data^5,6^. Integrating entropic measurements into structure based drug design will help enable the deliberate design of ligands based on total binding free energy by explicitly considering both enthalpy and entropy.

We previously showed that ligand binding redistributes protein entropy, with binding-site residues often becoming more rigid, while distal residues gain entropy, an effect estimated to contribute favorably up to ∼2 kcal/mol to the binding free energy^7^. Other studies have also observed similar patterns of redistributed entropy upon binding, suggesting that proteins often redistribute and fine tune conformational entropy to control function^8–11,12–149,15–17^. Yet, the molecular determinants by which ligands alter protein conformational entropy remain poorly understood. Our prior work indicated that specific interactions (i.e. hydrogen bonds) often decrease conformational entropy; however, substantial deviations from these trends highlight the underlying complexity of protein–ligand interactions^18^. Consistent with this, even minimal chemical changes, including the addition of a single methyl group, can produce strikingly different binding thermodynamics^17,19^. Beyond the protein, changes in solvent structure and dynamics are key to binding thermodynamics, with different systems demonstrating entropy being mainly derived from solvent or protein^20,21^. Ultimately, binding entropic effects emerges from the collective behavior of protein, solvent, and ligand^20,22^, supporting the joint examination of protein and solvent^23,24^.

To date, most insights on the entropic contributions to ligand binding come from molecular dynamics (MD) simulations and Nuclear Magnetic Resonance (NMR) relaxation measurements^9,15,16,25,26^. Yet both approaches are limited by low throughput, restricting their utility in high-throughput structure-based drug design. This contrasts sharply with X-ray crystallography, where throughput of data collection has rapidly improved due to multiple technological advancements, positioning it as the main technique for structure based structure drug discovery^27–29^. Although X-ray crystallography captures data from tens of thousands to billions of molecules with different conformations, due to ensemble averaging, most resulting models depict only a single static structure, leaving critical information about the multiplicity of states, and thus, entropy on the table.

Conformational heterogeneity, which is the energetic basis for protein conformational entropy, manifests as both anharmonic disorder (e.g., rotameric jumps) and harmonic disorder (e.g., small vibrations within a single rotameric well). While X-ray crystallography traditionally models disorder using B-factors (Atomic Displacement Parameters), these values often incorporate non-physical effects like modeling errors and crystal defects^30^.

Anharmonic motion is less commonly explicitly modeled, but can be done so with multiconformer or other ensemble modeling methods^31^. Accurately modeling both types of heterogeneity are essential for capturing protein conformational entropy^15,30,32,33^. To model both of these types of disorder, we utilize qFit to build multiconformer models^34,35^, which automatically models alternative conformers and B-factors as supported by the electron density. We have previously shown that these models can be used to examine how entropy may contribute to ligand binding^7^. The heterogeneity that drives protein conformational entropy is typically minute, often below an Angstrom, and distributed across the protein. Such subtle variations are readily masked by experimental noise or modeling bias^36^, underscoring the need for automated algorithms to detect and quantify them.

Here, we analyzed 100s of ligands bound to SARS-CoV-2 macrodomain (Mac1), an emerging antiviral target whose ADP-ribosylhydrolase activity counteracts the interferon response^37,38–42^. By treating crystallography as a statistical ensemble and analyzing its structural variation, we can disentangle authentic entropic signatures from noise^43,44^. We identified that in Mac1, similar protein-ligand binding interactions lead to similar magnitude and spatial distribution of protein conformational heterogeneity and solvent patterns. We connect these structural patterns with thermodynamic measurements of binding, revealing that increased protein conformational entropy upon binding and a less connected protein-solvent hydrogen bond network correlates with more entropic binding. Together, these findings and approaches lay the groundwork for incorporating conformational ensembles and water networks into structure-based drug design, enabling entropy to be leveraged prospectively.

## Results

### Dataset

From the UCSF QCRG AVIDD project, we curated a dataset of ligands bound to the SARS-CoV-2 Macrodomain (Mac1)^38,40,42,45^. We removed structures with ligands smaller than 200 Daltons or with occupancy less than 20% as estimated by PanDDA^46^. After these filtering steps, 316 structures remained, featuring ligands with molecular weights of 204.23–425.25 Da and occupancies ranging from 22% to 90% (**SFig. 1**). All structures were re-refined and then processed through qFit to obtain automated multiconformer models^34^, followed by a final round of refinement(**Methods; Supplementary Table 1**)^47^. As all liganded structures were obtained via compound soaking, the resulting models featured ligands with partial occupancies, introducing compositional heterogeneity between the bound and unbound structures. To correct for this, we developed occupancy-corrected bound models by comparing the ligand bound structure compared to apo structure, reweighting conformations based on ligand occupancy (**Methods**).

Using our occupancy-corrected models, we calculated crystallographic order parameters which combine the alternative conformers’ first torsion angle (χ1) and weighted B-factors, to estimated residue level conformatioanl heterogeneity. These were measured for all residues except glycine and proline(**Methods**)^48^. As an independent measure of conformational heterogeneity, we performed ensemble refinement for all structures to compute the change in root-mean-square fluctuation (ΔRMSF) for each residue (**Methods**)³⁰. Both of these metrics serve as proxies of changes in protein conformational heterogeneity representing protein conformational entropy.

### Ligand Binding Redistributes Conformational Heterogeneity to Three Intrinsic Regions

The apo and ligand-bound structures show no substantial backbone conformational differences, with the largest alpha carbon root mean squared distance (RMSD) being 0.16 Å (**SFig. 2A**). However, we identified that Mac1 structures tend to increase their conformational heterogeneity (referred here within as heterogeneity) mainly stemming from side chain fluctuations (**SFig. 2B-E**). We next asked where ligand binding alters the heterogeneity of Mac1. Most residues exhibited only minor shifts (**SFig. 3A**). However, a subset of residues consistently exhibited increased heterogeneity upon ligand binding (<-0.2 median Δorder parameter). These residues clustered in three localized regions: the loop–helix six segment beneath the distal ribose, the N-terminal end of helix 8, and helix 1(**SFig. 4**). To corroborate these findings, we analyzed the changes in RMSF determined with ensemble refinement. Residues that became more flexible based on RMSF (>0.25 median ΔRMSF), spatially overlapped with residues displaying increased heterogeneity. This suggests that upon ligand binding, Mac1 often redistributes protein conformational entropy to three distinct regions (**Figure 1A**).

**Figure 1.**
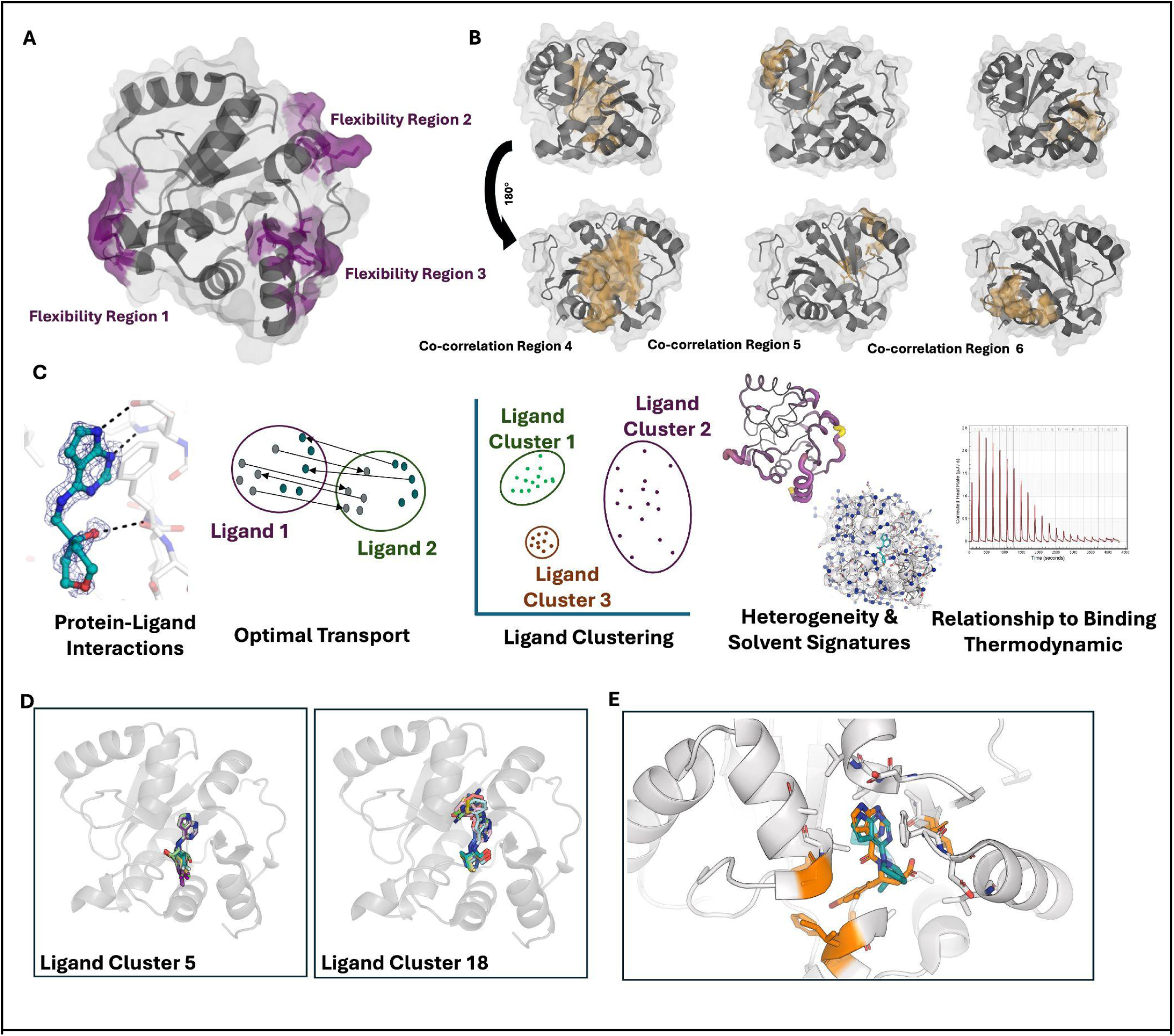
**A.** Residues that show a repeatable increase in flexibility upon ligand binding as measured by differences in order parameters or RMSF (region 1: residues 101-105, region 2: residue 157, 159, 161, 162, 165, region 3: 29, 30, 31). **B.** Residues with strongly correlated flexibility, revealing co-localized residues that change their conformational heterogeneity together (region 4: residues 36, 78-83, 93, 109, 110, 114-119, 142, 143, region 5: 37, 39, 45, 59, 60, 62, 94, 95, region 6: 13, 15, 16, 114, 146, 147, 148, 51, 52). **C.** Protein-ligand interaction fingerprints to relate to conformational and solvent signatures. All protein heavy atoms within 3 Å of any ligand heavy atom were used to define the binding interface. Graphs were compared using the fused Gromov–Wasserstein (FGW) distance, which jointly accounts for structural geometry and node feature differences. For each ligand cluster, we compare the conformational heterogeneity and solvent signatures and relate this to binding thermodynamics. **D.** Examples of ligands in cluster 5 and cluster 18, identifying different poses and binding site interactions. **E**. Protein-Ligand interfaces of cluster 10 (teal; PDB: 7HD6) v. 13 (orange; PDB: 7IJ0). Sticks represent binding site residues. Teal residues are only binding site residues for cluster 9, orange sticks are only binding site residues for cluster 18.

We analyzed apo structures to assess if the observed entropic redistribution reflects Mac1’s intrinsic properties. This analysis revealed that residues that gained heterogeneity upon binding were already significantly heterogeneous in the apo state. This suggests these intrinsically heterogeneous regions may act as entropic reservoirs, poised to undergo a significant change in conformational entropy upon ligand binding (**SFig. 5B**)^5^.

We next investigated if these flexible clusters act together by identifying residue clusters with a high correlation (Pearson coefficient >0.8) (**SFig. 6A**). We found no strong correlations within the most flexible regions. Instead, we identified three other residue regions that tend to change their flexibility together (**Figure 1B**), showing unit-like changes in ligand-modulated heterogeneity. Structural analysis revealed that these correlated regions are more rigid, have higher residue packing, contain more intra-protein hydrogen bonds, and are less solvent exposed than the highly heterogeneous regions (**SFig. 6B-D**). This suggests a division of heterogeneity upon ligand binding with the highly heterogeneous regions reflecting weakly constrained motions, which may exist in the apo structure and are used by the protein to ‘capture’ entropy, while the correlated regions display collective but constrained changes in heterogeneity. Overall, ligand binding alters both intrinsic entropic hotspots and collective, structurally connected regions.

### Ligand Properties Impact Protein Conformational Heterogeneity

Our next goal was to determine whether ligand properties determine how protein heterogeniety changes upon ligand binding. Consistent with our prior work^7^, ligands with higher molecular weight, fewer hydrogen bonds per molecular weight, higher logP, and more rotatable bonds tended to increase binding-site heterogeneity (**SFig. 7**). Across the full protein, only molecular weight correlated with overall protein heterogeneity (**SFig. 8**).

Given that many ligands in our dataset (n=52) exhibited alternative conformers, we next examined whether ligand properties were associated with alternative conformers. Ligands with alternative conformers tended to have higher molecular weight, fewer hydrogen bonds normalized by molecular weight, and higher logP values, indicating a lack of grounding interactions to the protein (**SFig 9**). Unsurprisingly, ligands with alternative conformers tended to increase conformational heterogeneity both within the binding site and across the protein (**SFig 10**).

### Graph-Based Optimal Transport Algorithm Identifies Protein–Ligand Interaction Clusters

We hypothesized that considering ligand properties in isolation would obscure how ligands alter protein heterogeneity, as such patterns likely emerge from the ways ligands interact with the protein^49^. Therefore, we developed a new graph-based algorithm to move beyond ligand properties alone. Our algorithm represents protein–ligand interactions as graphs resulting in “protein-ligand interactions” are represented by two matrices: a geometry matrix encoding pairwise interatomic distances, and a feature matrix containing chemical and structural values. We then apply the fused Gromov–Wasserstein optimal transport algorithm to capture both differences in interactions and chemical features^50^. We then identified ligand groups by clustering the squared Gromov–Wasserstein distance metrics with HDBScan (**Figure 1C; SFig. 11A**)^51^. This produced 25 clusters (3–21 structures each), with a median size of 4 structures per cluster (**Supplementary Table 2**). Ligand sizes vary moderately across clusters, with the largest average ligands in cluster 14 (median=28 atoms) and the smallest in cluster 38 (median=16 atoms).

We compared Tanimoto similarities within versus between ligand clusters and found that ligands within a cluster were substantially more similar to each other (median 0.50) than ligands across clusters (median 0.24; **SFig. 11B**). We also examined ligand properties across clusters but found no consistent trends (**SFig. 12**). Ligands modeled with alternative conformers tended to clustered together. We initially attributed the results to additional graph nodes from alternative conformers, but removing the ligand’s second conformation did not change the ligand clusters. Instead, higher Tanimoto similarity of ligands with alternative conformations, along with their non-grounding properties, suggests that these alternative states are enabled by shared structural features and weak anchoring interactions (**SFig. 13**).

Next we turned to how the ligand was interacting with the protein. All ligands tend to engage near space occupied by the adenosine group of ADPr in the product bound structure, with some clusters extending towards the first helix (ligand clusters 6, 14, and 15), showing a stronger interaction with residues 156, 22, and 52 (**Figure 1D/E**). In contrast, ligand clusters 2, 6, and 12 tended to have more engagement with residues 129-132, reaching slightly closer to the ADPr ribose site (**SFig. 14A/15**). We also identified hydrogen bond differences between the ligand clusters to residues 22, 126, and 157 (**SFig. 14B**).

### Protein-Ligand Interactions Drive Protein Conformational Heterogeneity Signatures

We then examined whether the different ligand clusters caused distinct changes in protein heterogeneity upon binding. We observed that different ligand clusters caused significantly different amounts of change in the protein’s overall heterogeneity (Kruskal–Wallis, p = 0.000152; **Figure 2A**). While we also noted significant changes within the binding site, clusters promoting increased heterogeneity across the entire protein did not always correlate with increased heterogeneity within the binding site (**SFig. 16A**). Since no single ligand property explained the large changes in protein heterogeneity, we propose that combining protein-ligand interaction data with ligand properties, as done with our optimal transport algorithm, enabled us to more accurately classify how ligands impact the entire protein, not just the binding site. We next examined whether other structural features tracked with changes in heterogeneity. We identify that ligand clusters with increased heterogeneity upon binding were less densely packed and formed fewer inter-protein hydrogen bonds (**SFig. 16B/C**), potentially points to how ligands drive changes in heterogeneity.

**Figure 2.**
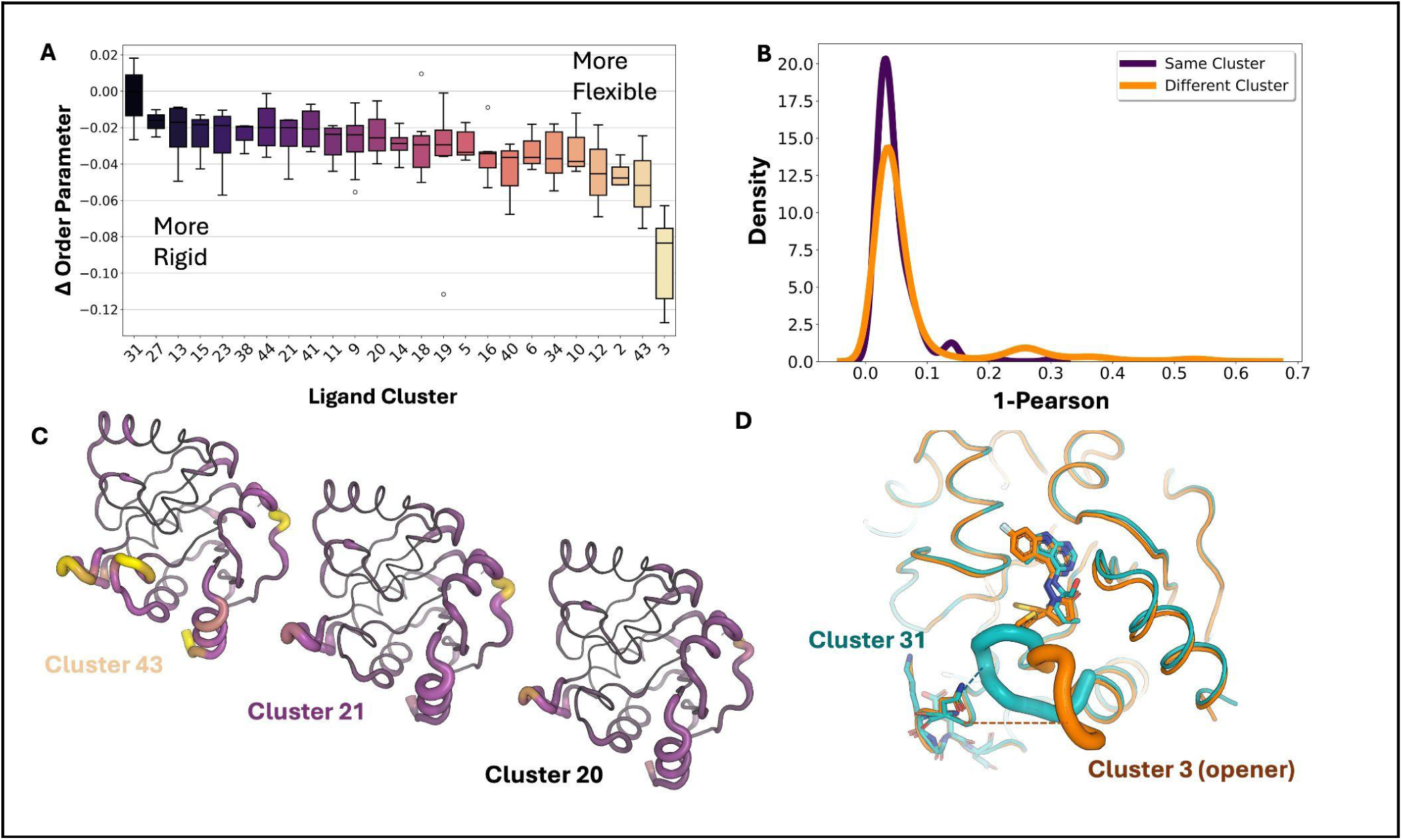
**A.** Change in overall estimated protein conformational heterogeneity upon binding across all PDBs in different ligand clusters. **B.** Lower (1–Pearson) values indicate higher similarity in the Euclidean distance of residue-level flexibility (order parameters); *p* = 2.6 × 10^−06^; Mann-Whitney. **C.** Differences in flexibility between ligand cluster 43, ligand cluster 21, and ligand cluster 20. Flexibility is shown with putty (larger = more flexible) and with colors (yellow = most flexible, black = most rigid). **D.** Example of ligand (orange) inducing an ‘opener’ (cluster 3; AVI-4200) conformation compared to a non-opener (cluster 31; AVI-4200) in teal. The opener loop is highlighted with residue 132 shown as sticks. The connection between the Phe132 loop and cluster 1 (residues 104-106) for both conformations is shown in dotted lines.

We then asked whether the different ligand clusters caused similar spatial changes in heterogeneity. Based on the Pearson correlation on the the delta order parameters of every residue, we found that ligands within the same cluster produced highly conserved spatial patterns of heterogeneity changes across the protein, meaning, that ligands in the same cluster tended to increase heterogeneity with the same set of residues (Mann–Whitney *p* = 2.6 × 10^−06^; permutation test, *p* = 0.002, *n* = 5,000; **Figure 2B/C**; **SFig. 16D**). These results show that distinct ligand–protein interaction networks leave specific spatial “signatures” of heterogeneity across the protein. This suggests that protein-ligand interactions, not just ligand properties, lead to magnitude and spatial differences in conformational heterogeneity.

Finally, we examined how the heterogeneity residue regions identified above changed across ligand clusters. Residue region one (residues 101–105) was the most heterogeneous and showed the largest variability across clusters (Kruskal–Wallis H-test; p=1e-7). The variability in this region mirrored global patterns of changes of heterogeneity, with ligand cluster 31 the least heterogeneous and cluster 3 the most (**SFig. 15**). To understand why heterogeneity in this residue region differed between ligand clusters, we computed residue interaction networks for each cluster. Ligand cluster 3 contained all previously identified “openers,” which shift Mac1 into a distinct state where the Phe132 loop adopts an open conformation^40^. This loop rearrangement disrupts its connectivity with residue region one, explaining why this region becomes substantially more heterogeneous in cluster 3 (**Figure 3D; Methods**).

**Figure 3.**
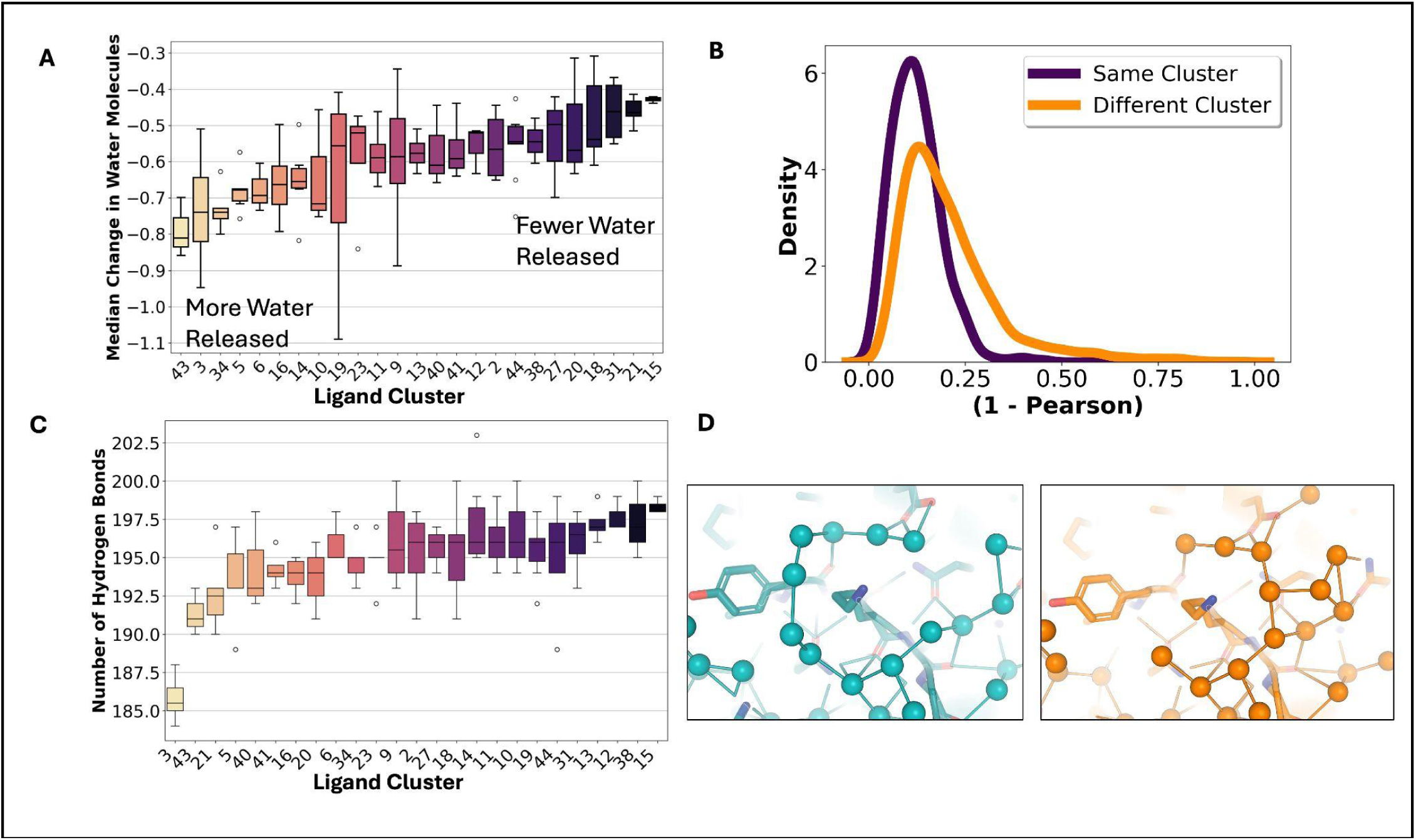
**A.** Across all structures, there was a loss in the number of water molecules bound, with ligand clusters varying in the number of water molecules released. **B.** Pearson similarity of the number of water molecules lost per residue in the same cluster versus in different clusters (*p* = 1.1e-37). **C.** Differences of the number of hydrogen bonds between protein, ligand, and water between different clusters. **D.** Differences in hydrogen bond networks in cluster 5 (PDB: 7HDC) versus cluster 40 (PDB:7HEN), demonstrating higher connectivity in cluster 38.

### Protein-Ligand Clusters Drive Solvent Signatures Across the Protein

Given prior evidence that ligand modifications can alter water-interacting networks, we next asked whether ligand clusters exhibited repeatable patterns of bound water molecules^52–54^. We began by examining the change in overall solvent exposure after ligand binding. We found that solvent exposure decreased across all ligand clusters, which likely results in the release of water molecules (**SFig. 18A**).

Our next step was to analyze changes in the modeled water molecules. Due to compositional heterogeneity, we could not distinguish between truly bound and unbound waters; however, we assumed that any observed differences largely reflected changes in the bound-state waters (as unbound waters should be consistent across clusters). To quantify this, we assigned each residue a “waters bound” score based on the number of water molecules found within a 3.2Å radius of its atoms (**Methods**). We then compared these counts for every residue between apo and all liganded structures. Most residues showed no change in water molecules (43.5%), followed by those that lost water molecules (41.7%), while only 14.8% of residues gained additional water molecules, mirroring the changes we see in the solvent accessible surface area. Ligand clusters that increased heterogeneity also tended to release more water molecules (**Figure 3A**). To observe how these water molecule changes were altered spatially, we compared the patterns of water molecules gained or lost across residues to see if they differed between ligand clusters. We found that the patterns of water loss/gain were significantly more correlated within ligand clusters than they were between different clusters (Mann–Whitney *p* = 1.1e-37; permutation test, *n* = 5,000; **Figure 3B/SFig. 18B**). We also looked at this trend with the binding site only, identifying the same trend (p = 1.25e-55, Mann-Whitney test; **SFig 18D**). These results indicate that more similar protein–ligand interactions give rise to more similar solvent organization throughout the protein.

We observed that ligand clusters which increased protein heterogeneity also corresponded to a higher number of released water molecules upon binding. Given this correlation, we hypothesized that these two phenomena may be functionally linked, especially in light of other data suggesting that proteins ‘borrow’ flexibility from surrounding water molecules^23,55^. We first looked at this from an individual residue perspective, however, we did not observe a strong relationship between residue heterogeneity changes and the number of water molecules gained or lost at that residue(**SFig 18C**). This suggested that local residue heterogeneity does not directly drive water release but instead may reflect a broader, coordinated system-level reorganization. Therefore, we hypothesized that clusters with fewer waters would show differences in their hydrogen-bonding networks between protein and solvent. We created graphs between all water molecules and all proteins, observing that clusters that had a higher flexibility upon ligand binding had fewer protein-solvent hydrogen bonds and lower overall connectivity of their hydrogen bond networks, potentially leading to these coupled changes (**Figure 4C/SFig. 19A; Methods**). Overall, this suggests a system-level reorganization of protein heterogeneity and the surrounding solvent.

**Figure 4.**
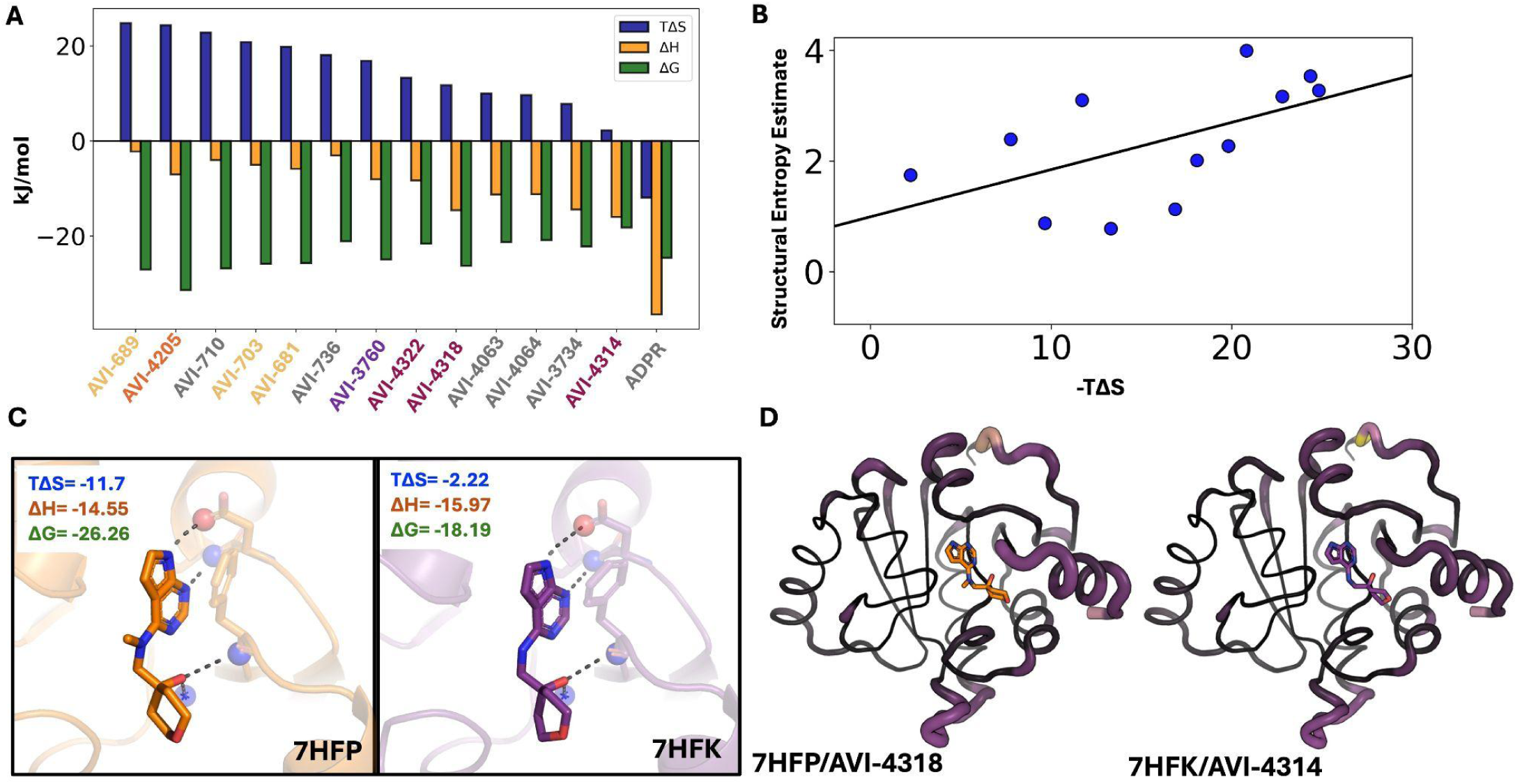
**A.** Thermodynamic binding values for ligands to Mac1. Ligands are colored based on their cluster (Cluster 3= yellow, Cluster 19 = orange, Cluster 11 = purple, Cluster 34 = maroon, No cluster = grey) **B.** Relationship between ITC-measured entropy and structural estimates of entropy (slope = 0.09, R² = 0.42) **C.** Structural differences in the binding site between PDBs 7HFP and 7HFK, which only differ by a single methyl group, yet have a 6 fold difference in entropy. **D.** Structural heterogeneity differences between PDBs 7HFP and 7HFK, demonstrating that 7HFK is more rigid upon ligand binding compared to 7HFP, partially contributing to entropy differences observed. Increase conformational heterogeneity is represented by thicker putty and light purple/yellow.

### Structural estimates of protein entropy correlate with ITC binding entropy

Previous studies have shown that protein conformational entropy, estimated from NMR relaxation order parameters, correlates with binding entropy^15,56^. However, it is unclear if crystallographic metrics could provide similar proxies. While crystallization can restrict certain motions present in solution, most heterogeneity arises from side chains, which remain capable of fluctuating within the crystal lattice. Additionally, X-ray crystallography measurements are not limited by the pico-nanosecond timescale of relaxation experiments. To test this, we performed ITC on a subset of ligands (n = 14), along with Mac1’s natural ligand ADPr (**SFig. 18/19; Supplementary Table 3; METHODS**). All ligands had at least some favorable entropic contribution to the free energy of association, in sharp contrast to ADPr, where entropy opposed binding (**Figure 4A**). Across all structures, we observed increased protein heterogeneity and a release of water molecules, potentially explaining why all ligands were shown to be entropically favorable. Notably, many of the most entropically favored ligands all come from ligand cluster 3, which all have alternative ligand conformations, increased protein heterogeneity, and fewer bound waters. All of these features are consistent with entropically favorable binding, underscoring the potential additive nature of the structural origins of entropy in protein–ligand interactions.

Next, we assessed whether protein entropy proxies obtained from the ensemble correlated with binding entropy, as has been shown for NMR relaxation experiments^15,56^. Protein conformational entropy was estimated using crystallographic order parameters, calculated by multiplying the average change in conformational entropy per residue by the total number of residues. While assuming residue independence overestimates the partition function, this approach provides a first-order approximation of protein conformational entropy. Solvent entropy was estimated from changes in polar and apolar solvent-accessible surface areas as previously described (**Methods**)^15,56^. The sum of protein, solvent, and ligand rotational/translation entropy is positively correlated with the entropic binding component measured by ITC (**Figure 4B**; slope = 0.09, R² = 0.42). While this is only a single system and the trend is weaker than what has been observed with NMR relaxation experiments, it provides an initial link between structural measurements of entropy from X-ray ensemble models and thermodynamic binding parameters.

We next examined how ligand properties influence binding thermodynamics. No ligand property significantly correlated with affinity, though molecular weight showed a slight positive trend (**SFig. 20**). In contrast, total binding entropy revealed clearer relationships: higher molecular-weight ligands exhibited significantly more entropy-driven binding (R² = 0.40, *p* = 0.015; **SFig. 21**). Ligands with higher logP values likewise tended to bind more entropically, as did those forming fewer hydrogen bonds per unit molecular weight, but neither was significant (**SFig. 22**). These thermodynamic trends parallel previous findings^57^ as well as our structural observations, with the properties associated with more entropic binding also being associated with increased conformational heterogeneity measured from structures, reinforcing the idea that ensemble-level structural features can illuminate the drivers of binding free energy. Notably, ligands modeled with alternative conformers were also more frequently associated with entropy-driven binding. However, because all ligands with alternative conformers and ITC measurements belong to cluster 3 (“openers”), it remains unclear whether this reflects a shared chemical scaffold, multiple ligand conformations, increased protein conformational entropy, or solvent release.

Among the ligands measured by ITC, we had multiple ligands from the same ligand clusters (3 and 34). Ligands in cluster 3 did not show much difference in the binding entropy across ligands. In contrast, the ligands in cluster 34 displayed more than a sevenfold difference in binding entropy. Within ligand cluster 34, PDBs 7HFP and 7HFN differ only by a sulfur-to-nitrogen substitution in their substituent group and exhibit similar entropic contributions (−11.7 vs. −13.3 kJ/mol). In contrast, 7HFK, which only differs from 7HFP by a single methyl group, shows a dramatically reduced entropic contribution of only −2.2 kJ/mol. Notably, the enthalpic components for 7HFP and 7HFK are nearly identical (−14.55 vs. −15.97 kJ/mol). Structural analysis revealed that 7HFP and 7HFK form nearly identical hydrogen bonds with the terminal carboxylate of Asp22, the backbone amides of Phe156 and Ile23, and a trapped water molecule, consistent with their similar binding enthalpies (**Figure 4C**). This suggests that entropy is the key differentiator in their binding affinities.

To understand why 7HFK exhibits lower binding entropy, we first examined changes in protein conformational entropy upon ligand binding. Although 7HFK shows a modest reduction in conformational entropy relative to 7HFP, contributing approximately ∼4 kJ/mol, this difference alone cannot account for the entropy discrepancy (**Figure 4D**). We next compared water structure around the two complexes and found that both bind nearly identical numbers of ordered water molecules (7HFP: −0.72; 7HFK: −0.74), suggesting water count is not a significant driver. Instead, differences emerged when analyzing the protein–water hydrogen-bond network. 7HFK forms a more connected and denser hydrogen bonding network. This strengthened hydrogen-bond network likely contributes to the less favorable binding entropy observed for 7HFK. From the structural ensemble models, we infer that the reduction in binding entropy in 7HFK is driven by a smaller increase in protein entropy and more rigid protein–solvent interactions. These findings mirror trends across the full dataset, where conformational heterogeneity and solvent interconnectedness covary, highlighting how subtle structural perturbations propagate through protein and solvent networks to influence entropic binding.

## DISCUSSION

Structure-based drug design has long prioritized static protein–ligand interactions while overlooking dynamic, entropy-driven forces that contribute nearly half of the binding free energy. Because protein conformational entropy is rarely quantified, its contributions, and the mechanisms that give rise to it, have remained largely unknown. Understanding how ligand binding reshapes conformational entropy and solvent networks, and how these changes can be harnessed for drug discovery, is still in its early stages^58^, but multiple studies have demonstrated the importance of considering entropy for affinity, selectivity, and cooperativity^7,10,17,19,58,59^, highlighting the need to systematically consider it. One of the largest challenges to quantifying entropy is that most observed changes are subtle, such as small shifts in side chain conformational heterogeneity, making it challenging to differentiate signal from noise. Approaching this problem through the statistical analysis of structural variation across hundreds of ligand-bound Mac1 structures, we were able to bring to disentangled entropic signals from noise^43,44^. We revealed how ligand binding reshapes conformational entropy and solvent networks, demonstrating how these signatures parallel experimentally measured binding entropy. Together, these results establish an initial framework for considering protein conformational entropy as tunable determinants of binding thermodynamics, advancing ensemble-based approaches to drug discovery.

To uncover how different ligands drive distinct entropic signatures, we modeled protein–ligand interactions as graphs and compared them using optimal transport. This enabled us to cluster ligands by both chemistry and interaction geometry. Ligands with more similar interactions produced similar changes in protein conformational entropy, both in magnitude and location, revealing that protein-ligand interactions propagate through the protein in reproducible ways. We also observed more similar changes in solvent networks among similar ligands, indicating that comparable interaction patterns reorganize bound water in consistent ways^60^. When only considering ligand chemistry, while we observed trends in the binding site in the change in magnitude of conformational entropy, similar to ligand properties known to correlate with more entropic binding^61^, we did not see trends across the protein, indicating that uncovering entropy signatures requires considering protein-ligand interactions.

Beyond identifying patterns of conformational entropy, we demonstrate that crystallographic proxies of conformational entropy carry quantitative thermodynamic meaning. Ligands that induced greater conformational entropy in the bound state had more entropically driven binding. We also observed that ligands with more entropic binding tended to displace more bound waters and formed fewer hydrogen bonds between proteins and solvent. These two reinforcing entropic mechanisms mirror patterns previously observed in catalysis^62,63^. While only demonstrated in a single protein, this relationship shows that protein conformational entropy inferred from crystallographic ensembles may be able to serve as a proxy for protein conformational entropy, analogous to the NMR-based “entropy meter”^15^.

We also showed that structural data can help identify why small changes between ligands can lead to drastically drastically different binding entropies. We observed that two ligands were only different by a single methyl group, yet have an almost seven-fold difference in binding entropy. This data is supported by the structural information, with changes in conformational entropy and hydrogen bonding patterns tracking with changes in binding entropy. We infer that this change in binding entropy is driven jointly by protein entropy and more rigid protein-solvent interactions. However, this still leaves open the important question of how to prospectively design to induce these favorable entropic changes.

Several important caveats should be considered when interpreting these findings. First, the partial occupancy of ligands necessitates careful deconvolution of conformational heterogeneity from compositional heterogeneity, which we addressed with our occupancy-corrected models, however, this potentially left mixed information on the bound and unbound state. With the growth of large-scale fragment-screening efforts, extracting meaningful conformational heterogeneity from data that is intermixed with compositional heterogeneity has become a major modeling challenge^64,65^. Automated modeling tools, such as qFit and PanDDA^34,46^, both used in this study, are starting points for modeling conformational ensembles, but further algorithmic development is needed to effectively tease out the role of protein conformational entropy in these studies. Further, there is a significant need for improvements in modeling water molecules, especially those that are partially occupied and intertwined with alternative side-chain states^66,67^. The importance of this is highlighted by the interdependence of proteins and solvent we observe, reflecting the fundamentally non-additive nature of binding entropy and affinity. Beyond this, the quantitative metrics of solvent and protein entropy from structures are still underdeveloped, especially in light of them not considering interdependence between states. Finally, we note that we attempted to apply a variety of machine learning frameworks to analyze this data, but it consistently led to overfitting. Despite the growth of protein–ligand datasets^68,69^, these datasets often remain too small to support conventional deep learning approaches. Future progress will depend on creative strategies, such as our optimal transport approach, that can extract robust patterns without overfitting, while simultaneously moving towards capitalizing on the expanding wealth of structural data with machine learning approaches.

Finally, one may question why we care about binding entropy at all. Molecular interactions driving affinity are inherently nonadditive; an intricate balance of attractive and repulsive forces that is nearly impossible to deconvolute^70,71^. Entropy is often described in terms of enthalpy–entropy compensation^72^. However, this relationship is not absolute; if it were, affinity could never be improved^1,61^. Designed ligands compared to natural ligands tend to be more entropically driven, likely due to the easier ability to increase affinity via entropy by increasing molecular weight or non-specific interactions (increased lipophilicity)^61^ and analyzes have shown these properties are associated with a higher probability of failure throughout clinical development^73,74^. However, we demonstrate that these poor ligand properties are not the only ways to entropically driven binding. Future studies with more careful control of these properties to determine if there are other ways to optimize binding entropy is needed. Finally, while different studies have demonstrated more beneficial enthalpic or entropic driven ligands^75,76^, ultimately, the best affinity ligands will come from optimizing for both thermodynamic properties where possible.

But likely more importantly than affinity optimization, the spatial changes in protein conformational entropy can be taking advantage of the effect of the ligand. Previous studies have demonstrated a key role of protein conformational entropy in cooperativity and allostery^8,12,17,19,58^, pointing to the need to more systematically study the spatial impacts of conformational entropy, as we can think of it as a tool not only to optimize affinity, but to optimize for selectivity and allosteric impacts. Mapping out these functional changes will be key for making more specific ligands. Our link between structural signatures and thermodynamic outcomes establishes a path for developing entropy-aware structure based drug design strategies, potentially enabling the rational design of ligands with entropically optimized affinity, selectivity, and allosteric function.

## METHODS

### X-ray crystallography

Mac1 crystals (P43 construct, residues 3–169) were grown by sitting-drop vapor diffusion in 28% wt/vol 570 polyethylene glycol 3000 and 100 mM *N*-cyclohexyl-2-aminoethanesulfonic acid pH 9.5 as described previously^40,45^. Compounds prepared in DMSO (100 mM) were added to crystal drops using an Echo 650 acoustic dispenser (final concentration of 10 mM)^77^. Crystals were incubated at room temperature for 2–4 hr prior to vitrification in liquid nitrogen without additional cryoprotection. X-ray diffraction data were collected at the Advanced Light Source (ALS beamline 8.3.1) or the Stanford Synchrotron Light Source (SSRL beamline 9–2). Data were indexed, integrated, and scaled with XDS^78^ and merged with Aimless^79^. The P43 Mac1 crystals contain two copies of the protein in the asymmetric unit (chains A and B). The active site of chain A is open; however chain B is blocked by a crystal contact. In addition, crystal packing in the chain A active site restricts movement of the Ala129–Gly134 loop, leading to decreased occupancy for compounds with substituents on the pyrrolidinone. To aid modeling the resulting conformational and compositional disorder, we used the PanDDA method^46^ to model ligands where the occupancy was low (<25%) or where there was substantial disorder. Data collection settings and statistics are reported in **Supplementary file 4**.

### Chemical synthesis

Unless otherwise noted, all chemical reagents and solvents used are commercially available. Chemicals represented with Z* were ordered from Enamine. The synthesis of the majority of other compounds are described in other papers^40,42,80,81^.

### Precursor molecules

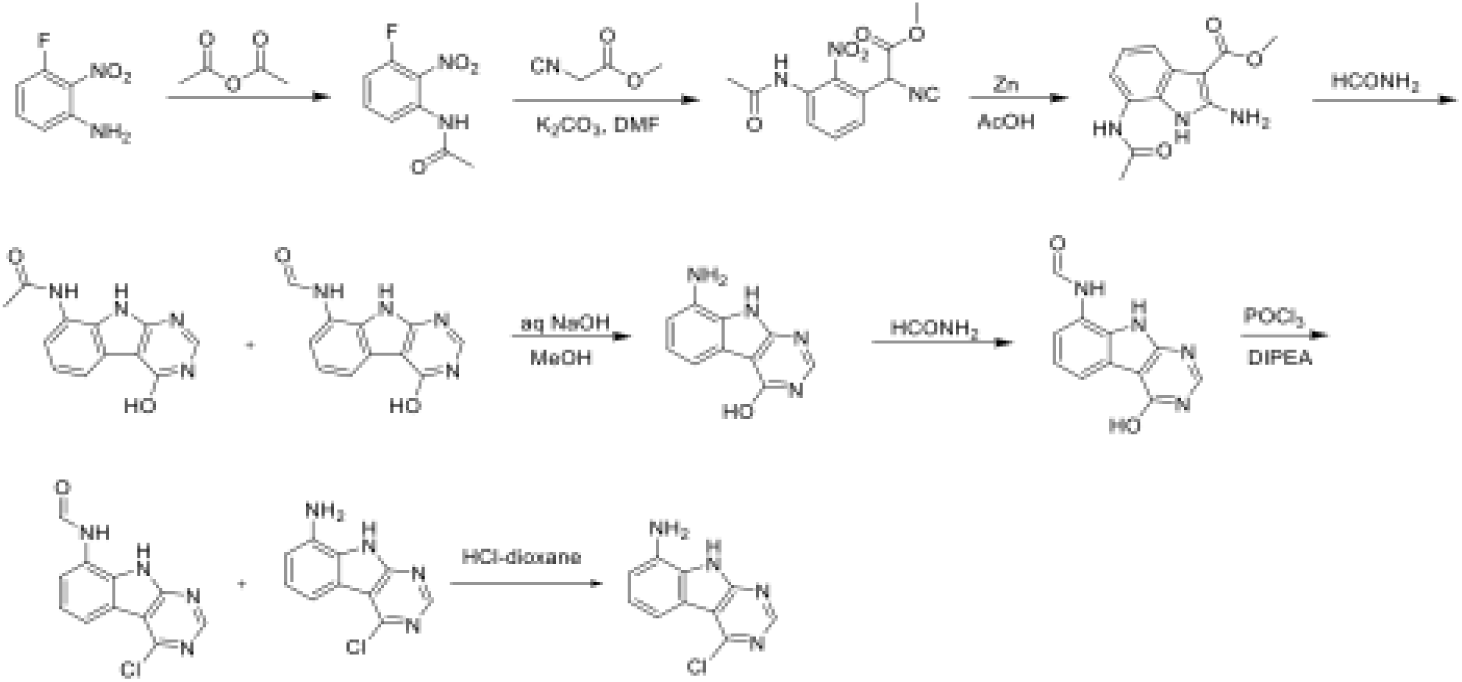

A solution 3-fluoro-2-nitroaniline (11 g, 70.51 mmol) in acetic anhydride (20 mL) was stirred at room temperature for 16 hours. The reaction mixture was filtered and the solids were washed with petroleum ether (100 ml) and dried to obtain 10.7 g (77%) of N-(3-fluoro-2-nitrophenyl)acetamide as a brown solid. LCMS (ESI): m/z= 199.3 (M+H)^+^

To a solution of N-(3-fluoro-2-nitrophenyl)acetamide (10.7 g, 54.04 mmol) in DMF (100 mL) was added methyl 2-isocyanoacetate (8.02 g, 81.06 mmol) and potassium carbonate (14.92 g, 108.08 mmol). After stirring at 80°C for 2 hours, the reaction mixture was cooled to room temperature, acidified with 2N HCl (ca. 2000 mL), and extracted with ethyl acetate (300 mL *3). The combined organic layers were washed with brine (100 mL), dried over Sodium sulfate and concentrated under reduced pressure. The residue was purified by silica gel chromatography (10: 1 petroleum ether/ethyl acetate) to obtain 11 g (73%) of methyl 2-(3-acetamido-2-nitrophenyl)-2-isocyanoacetate as a yellow solid. LCMS (ESI): m/z= 278.2 (M+H)^+^

To a solution of methyl 2-(3-acetamido-2-nitrophenyl)-2-isocyanoacetate (11 g, 39.71 mmol) in *glacial* acetic acid (100 ml), was added slowly zinc dust (25.81 g, 397.10 mmol) in two portions. After stirring at 60°C for 2 h, the reaction mixture was cooled to room temperature, filtered and washed with THF. The filtrate was concentrated under reduced pressure and purified by silica gel chromatography (10:1 dichloromethane/methanol) to obtain 6.2 g (63%) of methyl 7-acetamido-2-amino-1H-indole-3-carboxylate as a yellow solid. LCMS (ESI): m/z=248.3 (M+H)^+^

A solution of methyl 7-acetamido-2-amino-1H-indole-3-carboxylate (6.2 g, 25.10 mmol) in formamide (450 mL) was stirred at 220°C for 2 hours. The reaction mixture was then cooled to room temperature and poured in 100 ml of water. The resulting mixture was allowed to stand for 15 min before the solids were collected by filtration, washed with water, and dried to obtain 4.1 g of a 1:2 mixture of N-(4-hydroxy-9H-pyrimido[4,5-b]indol-8-yl)acetamide and N-(4-hydroxy-9H-pyrimido[4,5-b]indol-8-yl)formamide. This mixture was taken in methanol (25 mL) and aqueous 12 N NaOH (25 ml). After stirring at 60°C for 16 h, the reaction mixture was then cooled to room temperature, concentrated under reduced pressure to remove methanol and the residue was poured into 100 mL of water. The resulting mixture was allowed to stand for 15 min before the solids were collected by filtration, washed with water, and dried to obtain 3.5 g (70%) of 8-amino-9H-pyrimido[4,5-b]indol-4-ol as a brown solid. LCMS (ESI): m/z=201.2 (M+H)^+^

A solution of 8-amino-9H-pyrimido[4,5-b]indol-4-ol (3.5g, 17.5mmol) in formamide (30 mL) was stirred at 150°C. After 6 h, the reaction mixture was cooled to room temperature and poured into water (200 ml). The resulting mixture was allowed to stand for 15 min before the solids were collected by filtration, washed with water, and dried to obtain 3.5 g (88%) of N-(4-hydroxy-9H-pyrimido[4,5-b]indol-8-yl)formamide as a brown solid. LCMS (ESI): m/z=229.2 (M+H)^+^

To a solution of N-(4-hydroxy-9H-pyrimido[4,5-b]indol-8-yl)formamide (3.5 g, 15.35 mmol) in phosphorous oxychloride (30 mL) was added N,N-diiisopropylethylamine (5.94 g, 46.05 mmol), After refluxing for 16 hours, the reaction mixture was cooled to room temperature, concentrated and poured into water (20 mL). The resulting solid was filtered to obtain 500 mg of a mixture of N-(4-chloro-9H-pyrimido[4,5-b]indol-8-yl)formamide and 4-chloro-9H-pyrimido[4,5-b]indol-8-amine as a black solid. This mixture was taken in 4 N HCl in dioxane (15 mL). After stirring at room temperature for 4 h, reaction mixture was concentrated under reduced pressure, the residue was adjusted to pH7 with aq.Na_2_CO_3_, and extracted with EA (3 x 30 mL). The organic layers was dried over Sodium sulfate, concentrated under reduced pressure and the residue was purified by reverse phase chromatography (water/acetonitrile /0.1% ammonium bicarbonate) to obtain 320 mg (10%) of 4-chloro-9H-pyrimido[4,5-b]indol-8-amine as a white solid. ^1^H NMR (500 MHz, DMSO) δ 12.42 (s, 1H), 8.74 (s, 1H), 7.58 (d, *J* = 7.8 Hz, 1H), 7.25 – 7.08 (m, 1H), 6.93 (d, *J* = 7.7 Hz, 1H), 5.76 (s, 2H). LCMS (ESI): m/z=219.2 (M+H)^+^

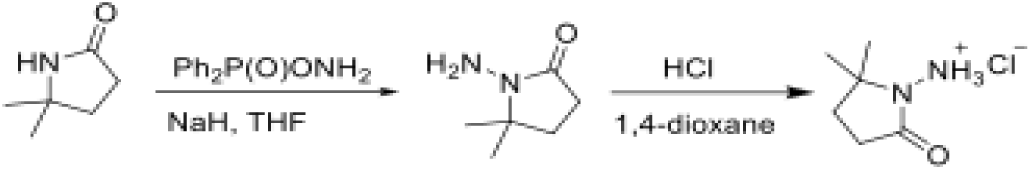

To a cooled (0°C) solution of 5,5-dimethylpyrrolidin-2-one (3 g, 26.54 mmol) in THF (60 mL) was added sodium hydride (2.13 g, 53.09 mmol), followed by addition of (aminooxy)diphenylphosphine oxide (12.4 g, 53.09 mmol) after 30 min. After stirring the resultant white suspension at 0C for 2 h, the reaction mixture was filtered through a Celite pad, the filtrate was concentrated and purified by silica gel chromatography (10:1 dichloromethane/methanol) to afford 3 g (75%) of 1-amino-5,5-dimethylpyrrolidin-2-one as yellow oil. LCMS (ESI): m/z= 129.1 (M+18)^+^;

A solution of 1-amino-5,5-dimethylpyrrolidin-2-one (1.5 g, crude) in 4 N HCl in dioxane (15 mL) was stirred at room temperature for 4h. The mixture was concentrated under reduced pressure, residue was triturated with diethyl ether and filtered to afford 1 g (53%) of 1-amino-5,5-dimethylpyrrolidin-2-one hydrochloride salt as a white solid. 1H NMR (500 MHz, DMSO) δ 9.48 (s, 3H), 2.39 (t, 2H, J = 7.8 Hz), 1.90 (t, 2H, J = 7.8 Hz), 1.30 (s, 6H). LCMS (ESI): m/z= 129.1 (M+18)^+^

### AVI-4205/RLA-5840 (RLA-T684B)

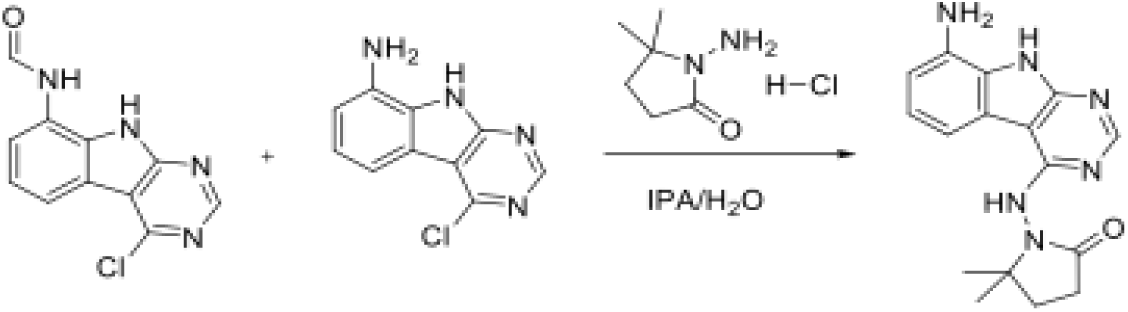

A solution of a mixture of N-(4-chloro-9H-pyrimido[4,5-b]indol-8-yl)formamide and 4-chloro-9H-pyrimido[4,5-b]indol-8-amine (15 g, 60.97 mmol) and 1-amino-5,5-dimethylpyrrolidin-2-one hydrochloride (11.7 g, 91.46 mmol) in isopropanol (150 mL) and water(15 mL) was stirred at 100°C for 16 hours. The reaction mixture was diluted with water (30 mL) and extracted with ethyl acetate (3 x 60 mL). The combined organic layers were dried over Sodium sulfate, concentrated under reduced pressure and the residue was purified by reverse phase chromatography (water/acetonitrile /0.1% ammonium bicarbonate) to obtain 77.3 mg (10.2%) of 1-((8-amino-9H-pyrimido[4,5-b]indol-4-yl)amino)-5,5-dimethylpyrrolidin-2-one as a white solid. 1H NMR (500 MHz, MeOD) δ 8.32 (s, 1H), 7.56 (d, 1H, J = 7.8 Hz), 7.12 (s, 1H), 6.85 (d, 1H, J = 7.6 Hz), 2.56 (t, 2H, J = 7.9 Hz), 2.17 (s, 2H), 1.37 (s, 6H). LCMS (ESI): m/z 311.3 (M+H)^+^.

### Creating multiconformer and ensemble models

All structures were modeled using the qFit pipeline (version 2024.2)^34,47,82^. Ligands were kept in their initially modeled conformation, including multiple conformations where modeled. qFit structures were then placed through Phenix.refine (version 1.20).

All structures were also run using ensemble refinement (version 1.20). We selected TLS (0.8), Wxray_coupled_tbath_offset (5.0), and tx (0.8) based on a grid search across apo mac1 (PDB: 7KQO) that provided the lowest Rfree. We then used these same parameters to run ensemble refinement on all structures. All ligand occupancies were kept as what was determined by the PanDDA algorithm and subsequent refinement optimizing the B-factor and occupancy of the ligand.

### Creating occupancy-corrected models

All structures shared the same unit cell and space group, each containing two protomers. We restricted our analysis to the A chain as in this crystal form, ligands bind only to the A chain, as the binding site in the B chain is blocked by a crystal contact. Since all ligands exhibited occupancies below 1, any observed conformational heterogeneity reflects a mixed population containing both the apo (unbound) and ligand-bound protein states.

For instance, a ligand occupancy of 50% means the experimental signal is derived equally from the ligand-bound and unbound conformations. Therefore, we developed an algorithm to correct the model to only account for the bound state, creating occupancy-adjusted models. To do this, we took the ligand structural multiconformer models derived using qFit, as well as a multiconformer model of the apo state (PDB: 7KQO). We also had the estimated ligand occupancy from PanDDA and subsequent optimization with refinement.

In multiconformer models, if residues are modeled in multiple conformations, each conformation is represented by coordinates, B-factors, and occupancy. The assumption behind the algorithm is if a ligand structure had 50% ligand occupancy and a residue was modeled with two conformations, each with 50% occupancy, one that matched the apo structure and one that did not, we assume that the ‘true’ bound state conformation of that residue would be the one not found in the apo structure. Therefore, our goal was for every residue, adjust the occupancy of ligand structure based on the ligand occupancy and the overlap of the ligand models compared to the apo model.

To reweight the occupancy of the residues, we took the apo structure (PDB: 7KQO) and a ligand structure and performed a local backbone alignment (5 residues) between the apo model and ligand model. We then compared the conformations of the ligand model to the apo model. If both models only had a single conformer for that residue, we did nothing. If the ligand model did not overlap with the apo model (determined by all chi angles being within 15**°**), we did nothing. However if any conformations of the ligand model did overlap with the apo model, we reweighted the conformations of the ligand model based on the ligand occupancy. For any overlapping conformation(s), we downweighted the conformation of the apo structure by 1-ligand occupancy. We then rescaled the occupancy of the other conformations of the residues.

### Order Parameter Calculation

To integrate the anharmonic fluctuations between alternative conformers with the harmonic fluctuations modeled by B-factors^30^, we used a crystallographic order parameter^48^. Order parameters provide information on each side chain’s first torsion angle (χ1) for all residues except glycine and proline. Order parameters are measured on a scale of 0–1, with 1 representing a fully rigid residue and 0 representing a fully flexible residue. These were measured on the occupancy-corrected multiconformer models.

### Root mean squared fluctuations (RMSF)

RMSF was chosen over root-mean-square deviation because many alternative conformers were predicted to have the same occupancy, making it difficult to define the main conformer. RMSF was measured for each residue based on all side-chain heavy atoms. RMSF finds the geometric center of each atom in all alternative conformers. It then takes the distance between the geometric mean of each conformer’s side-chain heavy atoms and the overall geometric center. It then takes the squared mean of all of those distances, weighted by occupancy.

### Sequence Conservation

To determine the conservation of residues, we used Consurf^83^ via their online portal. ConSurf estimates the evolutionary conservation of amino acid positions in a protein by analyzing multiple sequence alignments of homologous sequences. It assigns a conservation score to each residue based on the rate of evolutionary change, inferred from a phylogenetic tree using maximum likelihood, assigning each residue with a value between 0 (not conserved) to 9 (highly conserved). For each conservation category, we then examined the distribution of order parameters of those residues.

### Ligand Properties

Ligand properties were calculated using RDKit via python by inputting their SMILE strings.

### Optimal Transport Algorithm

We computed the structural similarity between protein binding pockets using the Fused Gromov–Wasserstein (FGW) distance^50^, which integrates both geometric and chemical information of nodes. We first built a graph of the ligand and all protein heavy atoms within 3 Å surrounding the ligand. We then represent all atoms included by two matrices: a geometry matrix (C) encoding pairwise interatomic distances, and a feature matrix (F), which include the following variables one hot encoded: element type, occupancy, residue type, and if it is a ligand or protein. These matrices describe both the spatial layout and atomic composition of the binding environment, while uniform weights assign equal probability to all atoms within the pocket. Pairwise comparisons between pockets are then performed using the fused Gromov–Wasserstein optimal transport framework implemented in the Python Optimal Transport (POT) library. This method computes the minimal transport cost required to align one pocket’s geometry and feature distribution with another’s, controlled by a balance parameter α (0.7) that weights geometric versus chemical contributions. The algorithm outputs a symmetric distance matrix, where smaller values indicate more similarity. This framework provides a quantitative, rotation- and translation-invariant measure of pocket similarity, allowing direct comparison of protein–ligand environments across multiple structures. We then used the FGW value to cluster all pairwise binding site comparisons with HDBSCAN^51^, enforcing the cluster to be at least 3 members.

### Residue Packing

We quantified the residue packing density of every residue. We first remove hydrogens and non-protein atoms. For every residue, we consider all alternative conformers and build a spatial index to find all nearby atoms in 3D space. We then quantify the packing of the residue by considering the number of residues with at least one heavy atom within 5 Å of any atom of the base residues. For residues with alternative conformers, we weighted this by the higher occupancy of two interacting proteins. We only considered residues in chain A when calculating residue packing.

### Hydrogen Bond

We first parse the PDB file identifying potential hydrogen bond donors and acceptors based on backbone and sidechain chemistry. For each residue, if there are no alternative conformers, we apply distance and angle cutoffs (default 3.5 Å donor–acceptor distance and 120° angular threshold) looking at neighboring residues and/or water molecules. If the residue has an alternative conformer, we only allow hydrogen bonds to alternative conformers of the same label (i.e. A altlocs with A altlocs). Hydrogen bonds are identified if these geometric criteria are satisfied, using either explicit hydrogen positions (on protein atoms) or approximated heavy-atom vectors (on water molecules, where hydrogens are missing). We only considered residues in chain A when calculating the hydrogen bonds. To calculate this with water, we considered chain A and water molecules with 4Å of any heavy atom in chain A.

The detected interactions are assembled into a network representation where residues (and subsequently water molecules) form nodes and hydrogen bonds form edges, annotated with bond counts, distances, angles, and interaction categories. From this graph, residue-level network properties such as degree and betweenness centrality are computed.

### Correlation

To detect correlated heterogeneity of residues, we calculated Pearson correlations of delta order parameters across all residues and structures. We then identify highly correlated residues as those with Pearson correlation >0.8.

### Comparing Protein-Ligand Interactions

To compare ligand interactions between clusters, we determined expected interactions (based on all protein-ligand interactions) versus observed interactions (within each ligand cluster). We defined any interaction if pi-pi stacking, van der Waals, or hydrogen bond was detected. A van der Waals contact was defined as any non-bonded heavy-atom pair with an interatomic distance ≤ 4.0 Å. π–π interactions were defined between aromatic rings when the centroid–centroid distance was ≤ 5.0 Å and the angle between ring normals ≤ 30°. A hydrogen bond was defined as a donor–acceptor heavy-atom distance ≤ 3.5 Å and a donor–H–acceptor angle ≥ 135°.

### Residue Interaction Network

To identify common residue interaction networks (RINs) from hydrogen-bond data across multiple structures in the same clusters, the script takes in the hydrogen bond analysis above and converts donor–acceptor pairs into residue-level undirected edges. Within each cluster, it aggregates these edges across all PDBs in each cluster, yielding a set of cluster-specific residue–residue connections. The algorithm then compares edge prevalence between clusters, computing odds ratios, prevalence differences, and Fisher’s exact p-values to identify connections that are statistically enriched or depleted in one cluster relative to the rest.

### Solvent-exposed surface area

We calculated solvent-exposed surface area (SASA) using FreeSasa^84^, getting SASA values for each atom. Polar SASA was defined as SASA for N, O, and S atoms, whereas apolar SASA was defined for C atoms. We used total SASA, rather than binary information, so we did not apply a relative solvent accessible surface area cut off.

### Water Counts

We quantified local solvation by counting the number of water molecules within 3.2 Å of each residue in the structure. For each PDB file, all non-hydrogen atoms of protein residues were compared against the oxygen atoms of water molecules, and any water whose oxygen atom lay within 3.2 Å of any heavy atom in a given residue was counted once. This analysis generated residue-level water contact counts, providing a quantitative measure of local hydration and solvent accessibility across the protein surface.

### Isothermal Titration Calorimetry

Mac1 samples were purified as previously described^38^; however, the final SEC buffer was 150mM NaCl and 20mM TrisHCl. The protein concentration was determined by UV absorption at 280 nm. The ligand stocks, except ADPr, were kept in DMSO stocks of 100mM. Ligand, from 100mM DMSO stocks, was dissolved in 3-5mM of the same SEC buffer as the protein. Protein stocks (all 4mM) were then matched with the same DMSO concentration as the ligand. ITC experiments were performed on Affinity ITC from TA Instruments at pH 7.0 and a temperature of 298 K by titrating the protein at a concentration of 4 mM into the cell containing the ligand at a concentration of 3-5uM. 24-26 injections were collected per experimental run, with the first injection being 1uL, the rest being 2uL. Peak integration was done using NITPIC^85^, with individual error bars assigned to each injection. We expected a single binding site, so measurements with an n more than 0.4 away from 1 were removed from subsequent analysis. A single-site binding model was simultaneously fitted to the titration curves to yield the binding enthalpy (ΔH°), the fraction of binding-competent protein (n), and the dissociation constant (Kd).

### Estimates of Entropy

Estimation of protein conformational entropy was determined by calculating the average difference in order parameters for every residue and multiplying this by the number of residues in chain A of the macrodomain (164 residues). Solvent entropy was calculated by looking the changes in apolar and polar surface area from apo to bound using the following formula:

Solvent Entropy = 0.45 * ΔSASAapolar - 0.26 * ΔSASApolar^86^

## Availability of Code

All analysis code is available at: https://github.com/Wankowicz-Lab/ensemble_bioinformatic_toolkit

qFit code and refinement scripts are available at: https://github.com/ExcitedStates/qfit-3.0/tree/main/src/qfit

## Acknowledgements

We thank Willow Coyote-Maestas for his helpful comments. Our work is supported by NIH U19AI171110 (JSF, ARR, & SAW) and by the American Cancer Society (SAW).

## Supplementary Figures

**Supplementary Figure 1.**
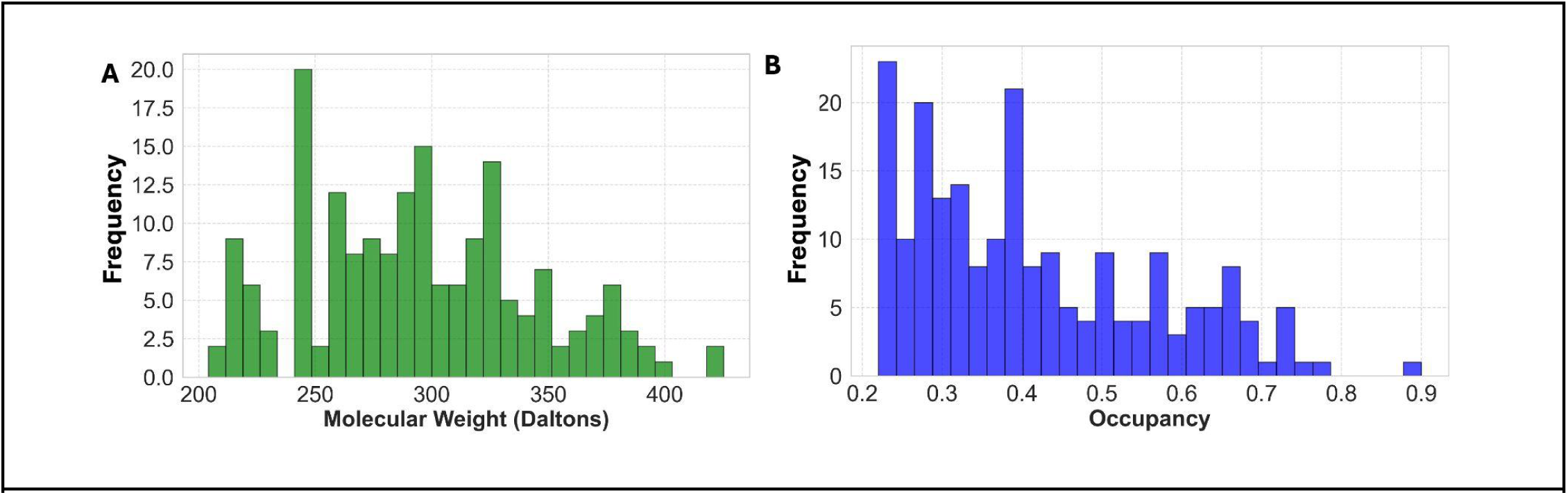
**A.** Distribution of molecular weight (range: 204.23-425.25). **B.** Distribution of ligand occupancy (22%-90%).

**Supplementary Figure 2.**
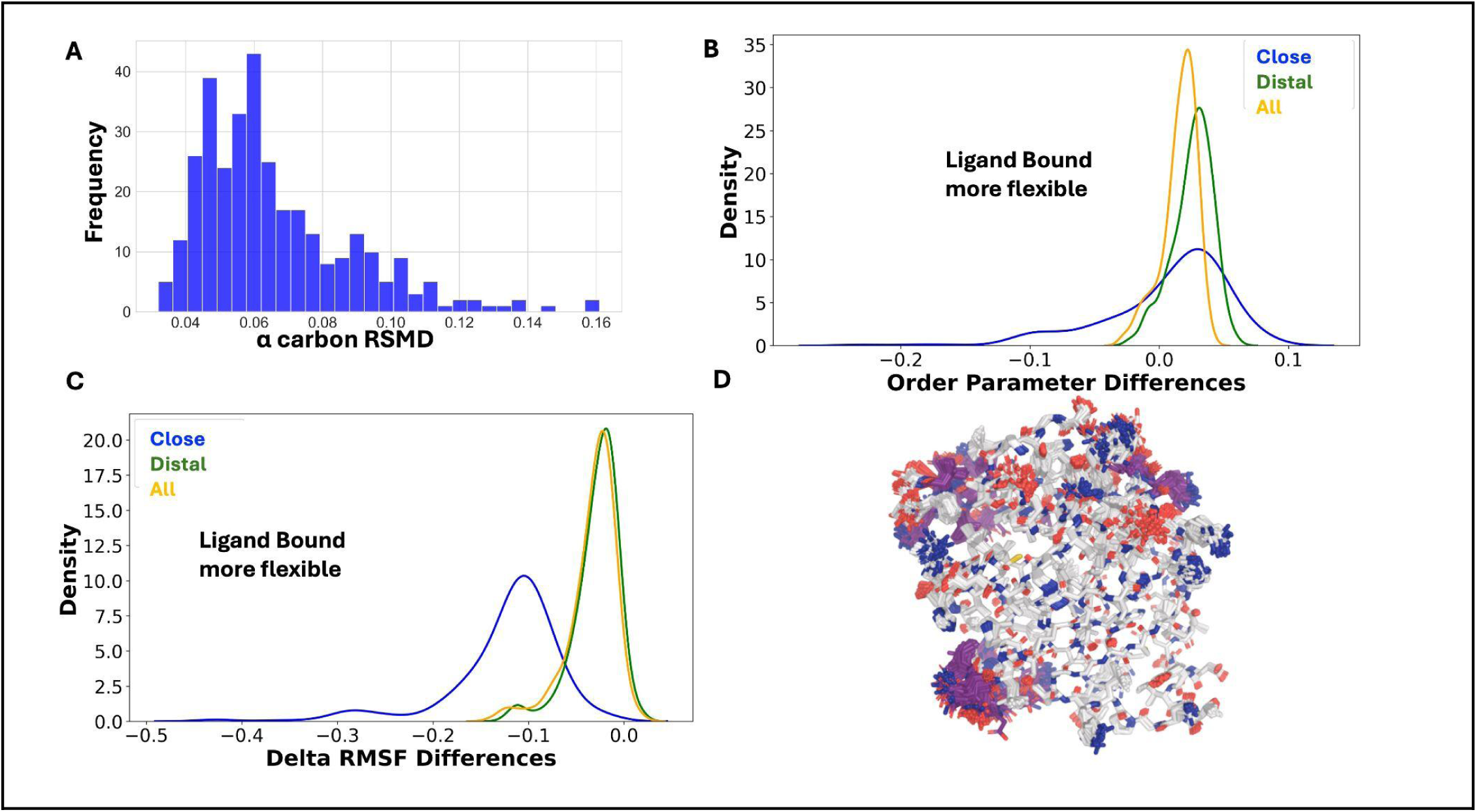
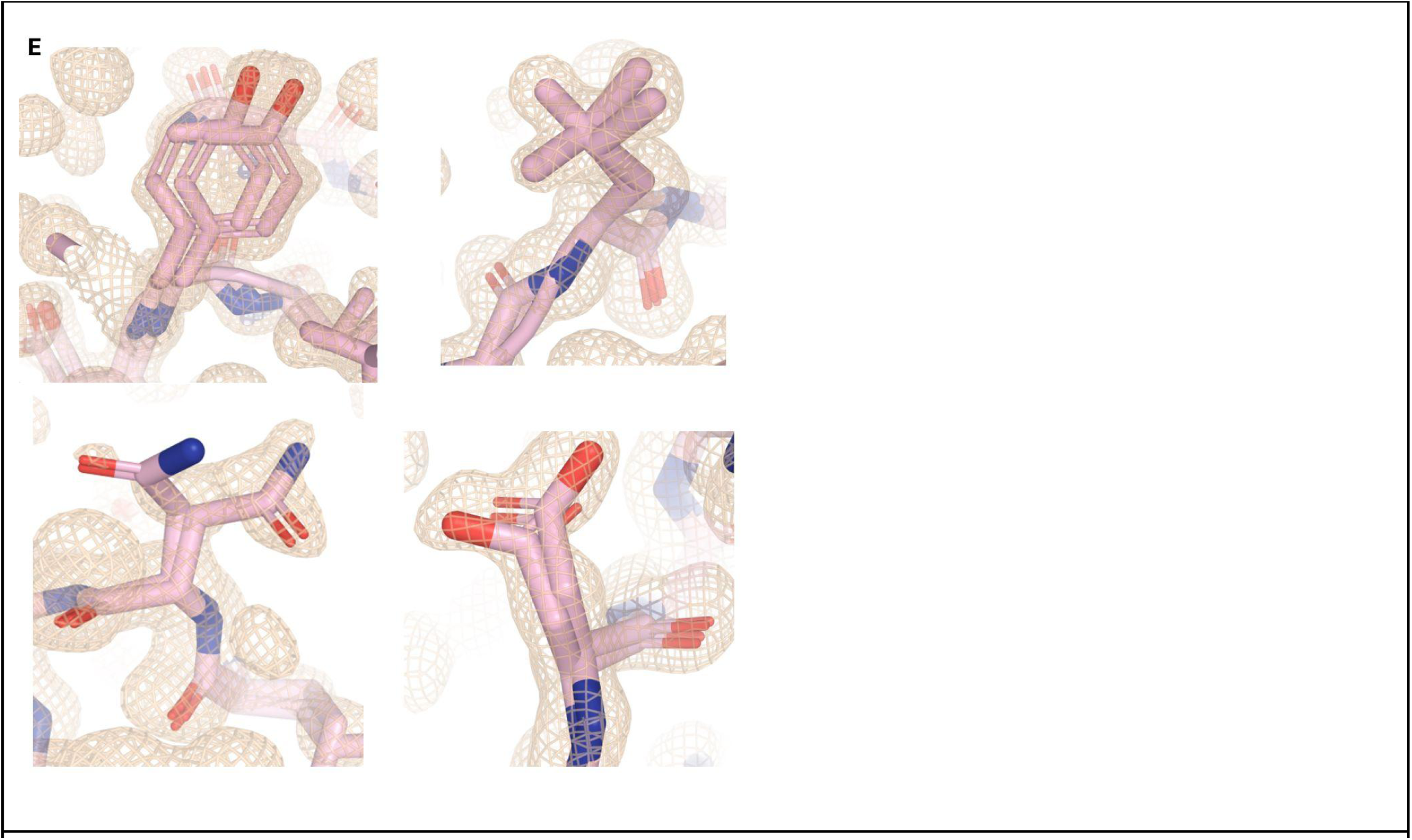
**A.** Distribution of the alpha carbon RMSD values between apo and ligand-bound structures of Mac1. While most structures exhibit minimal backbone deviation, the distribution shows a long tail, indicating a subset of structures with larger backbone conformational differences. **B.** The average delta order parameter across all ligand-bound structures separated by location. Binding site residues are defined as those within 4 Angstroms of any ligand heavy ligand and distal residues as those more than 10 Angstroms from any ligand heavy atom **C.** The average delta RMSF across all ligand-bound structures separated by location. **D.** Example of ensemble refinement output with flexible regions colored purple. **E.** Examples of multiconformer side chains identified by qFit and used to model the anharmonic heterogeneity present.

**Supplementary Figure 3.**
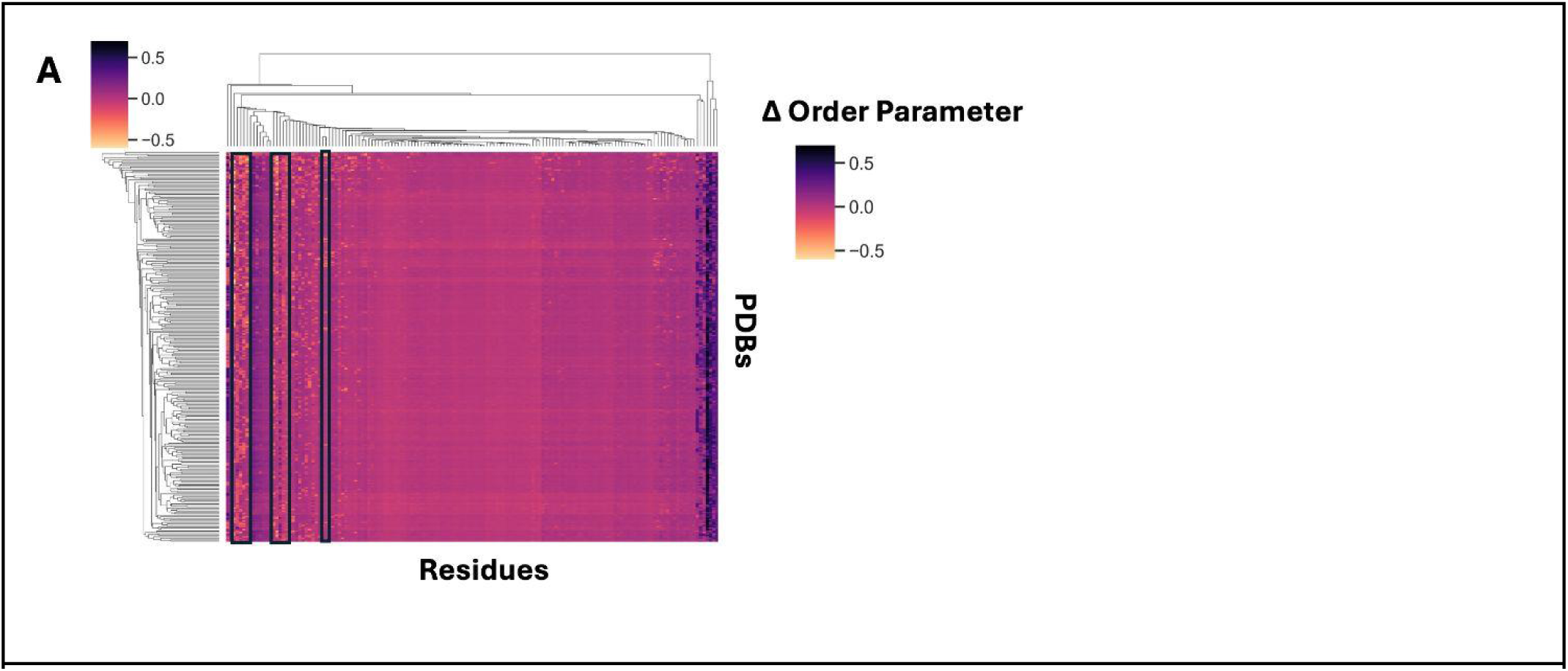
**A.** Heatmap of residues that gain or lose conformational heterogeneity upon ligand binding across all macrodomain structures. Dark purple indicates an increase in conformational heterogeneity upon binding.

**Supplementary Figure 4.**
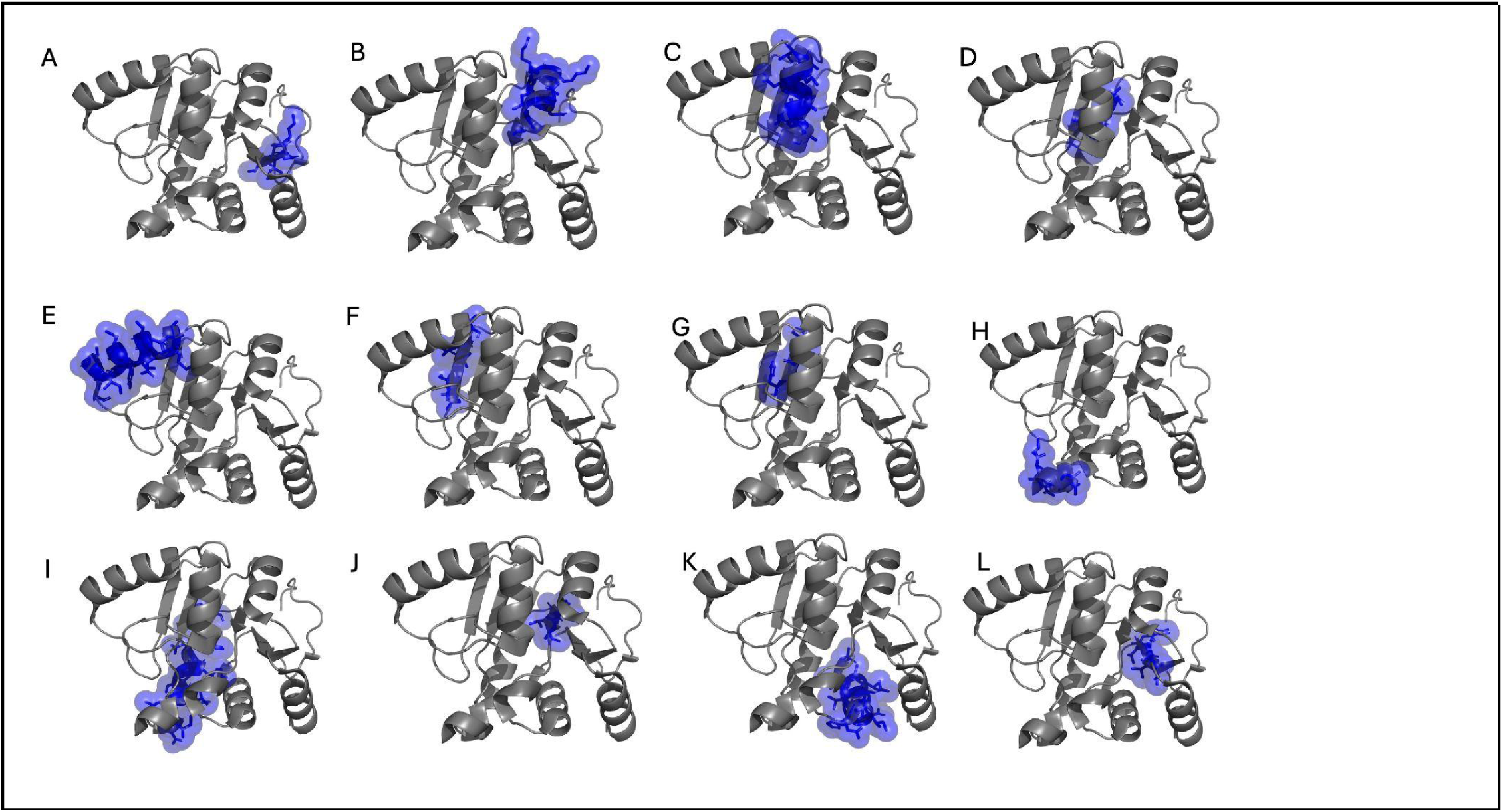
Mac1 Locations **A.** beta sheet 1, **B.** alpha helix 1, **C.** alpha helix 2, **D.** beta sheet 2, **E.** alpha helix 3, **F.** beta sheet 3, **G.** beta sheet 4, **H.** loop 100-105, **I.** alpha helix 4, **J.** beta sheet 5, **K.** alpha helix 5, **L.** beta sheet 6

**Supplementary Figure 5.**
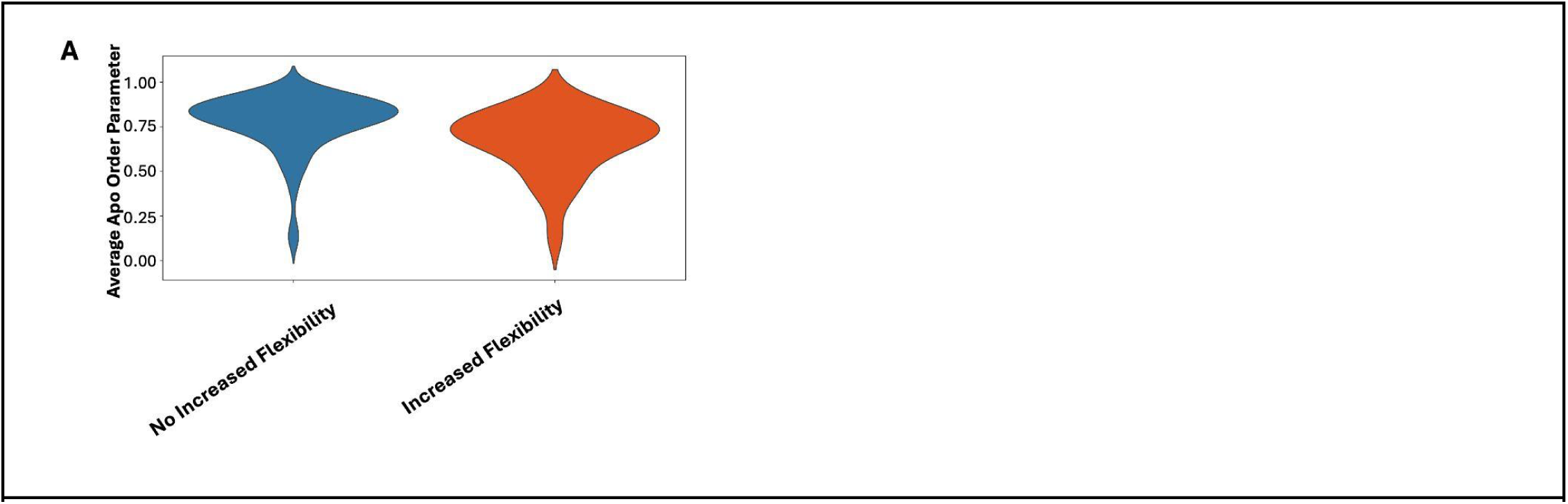
**A.** Average apo crystallographic order parameter between residues identified has having repeatable increase in flexibility upon ligand binding v. those without repeatable increase in flexibility.

**Supplementary Figure 6.**
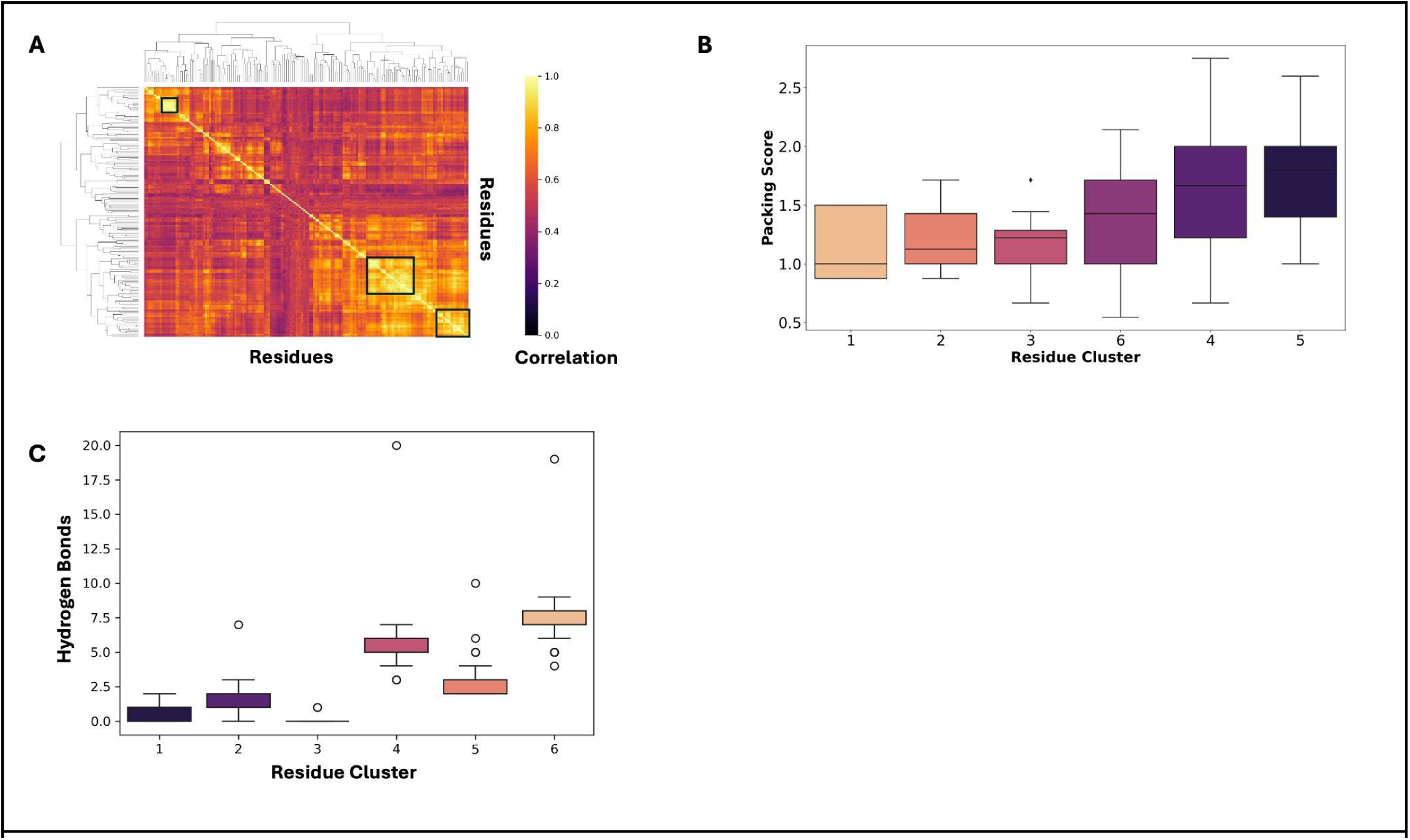
**A.** The correlation matrix of residue flexibility, where positive correlations (yellow) indicate that residues tend to fluctuate together. **B.** Distribution of packing per residue cluster. Residue clusters with overall increased flexibility tend to be less well packed (clusters 1,2,3). In contrast, residues clusters with high co-correlation tend to be well packed (clusters 4,5,6). **C.** Distribution of hydrogen bonds within cluster residues (normalized by number of residues).

**Supplementary Figure 7.**
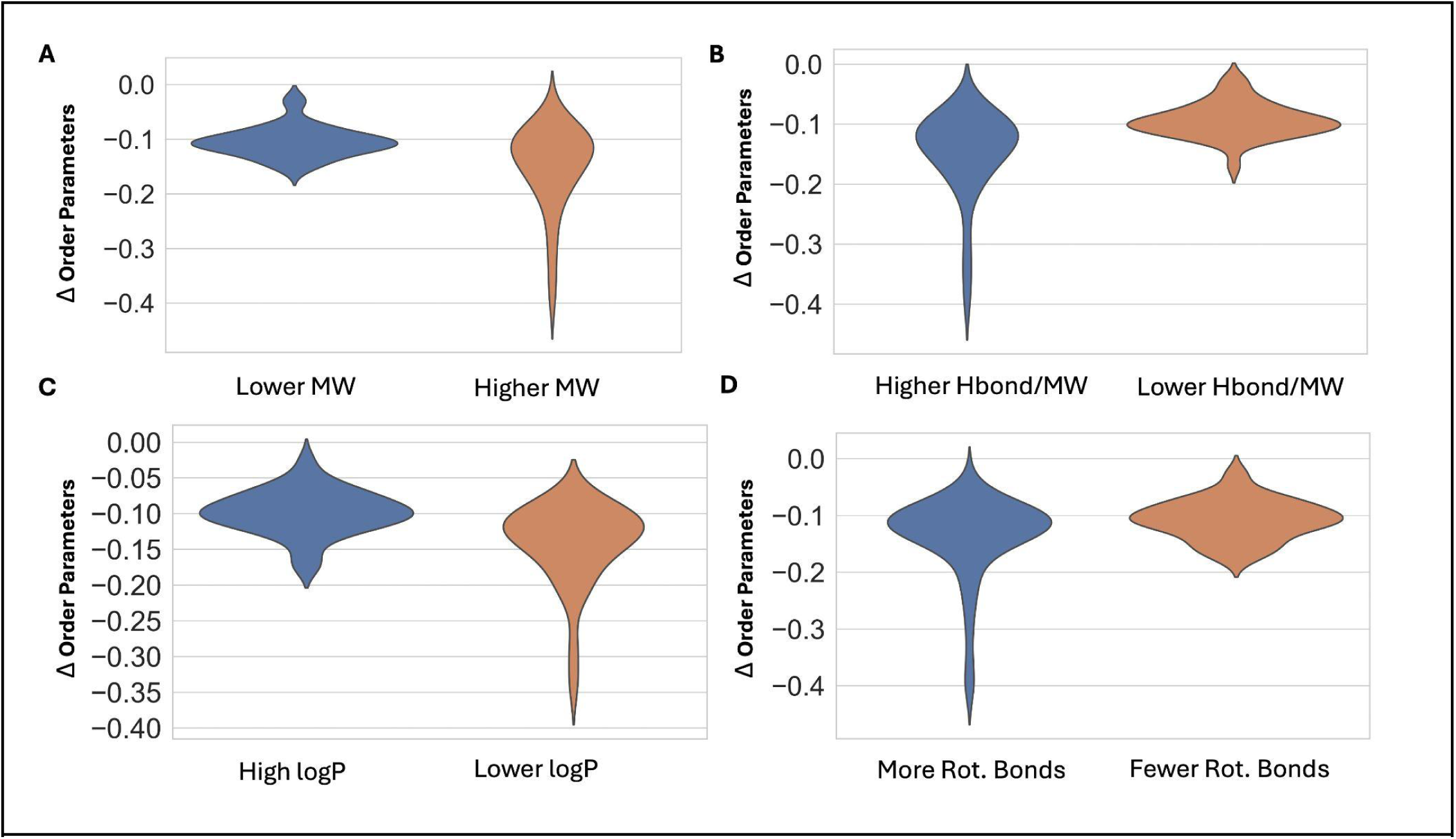
Ligand properties correlation with conformational heterogeneity within the binding site. **A.** Higher molecular weight is more flexible (Higher Molecular Weight: ≥343.63 Da; ΔOP −0.11, Lower Molecular Weight: ≤ 270.38; ΔOP −0.11; p=0.01; Mann Whitney U test), **B.** More hydrogen bonds/molecular weight is less flexible (More Hbond/MW ≥0.04; ΔOP −0.09, Less Hbond/MW: ≤0.3; ΔOP −0.12; p=3.2e-05), **C.** High logP is more flexible (Higher logP ≥0.27; ΔOP −0.09, Lower logP ≤-1.17; ΔOP −0.09; p=9.5e-05) **D.** Higher number of rotatable bonds is more flexible (More rotatable bonds ≥5; ΔOP −0.11, Fewer rotatable bonds ≤3; ΔOP −0.11; p=0.03).

**Supplementary Figure 8.**
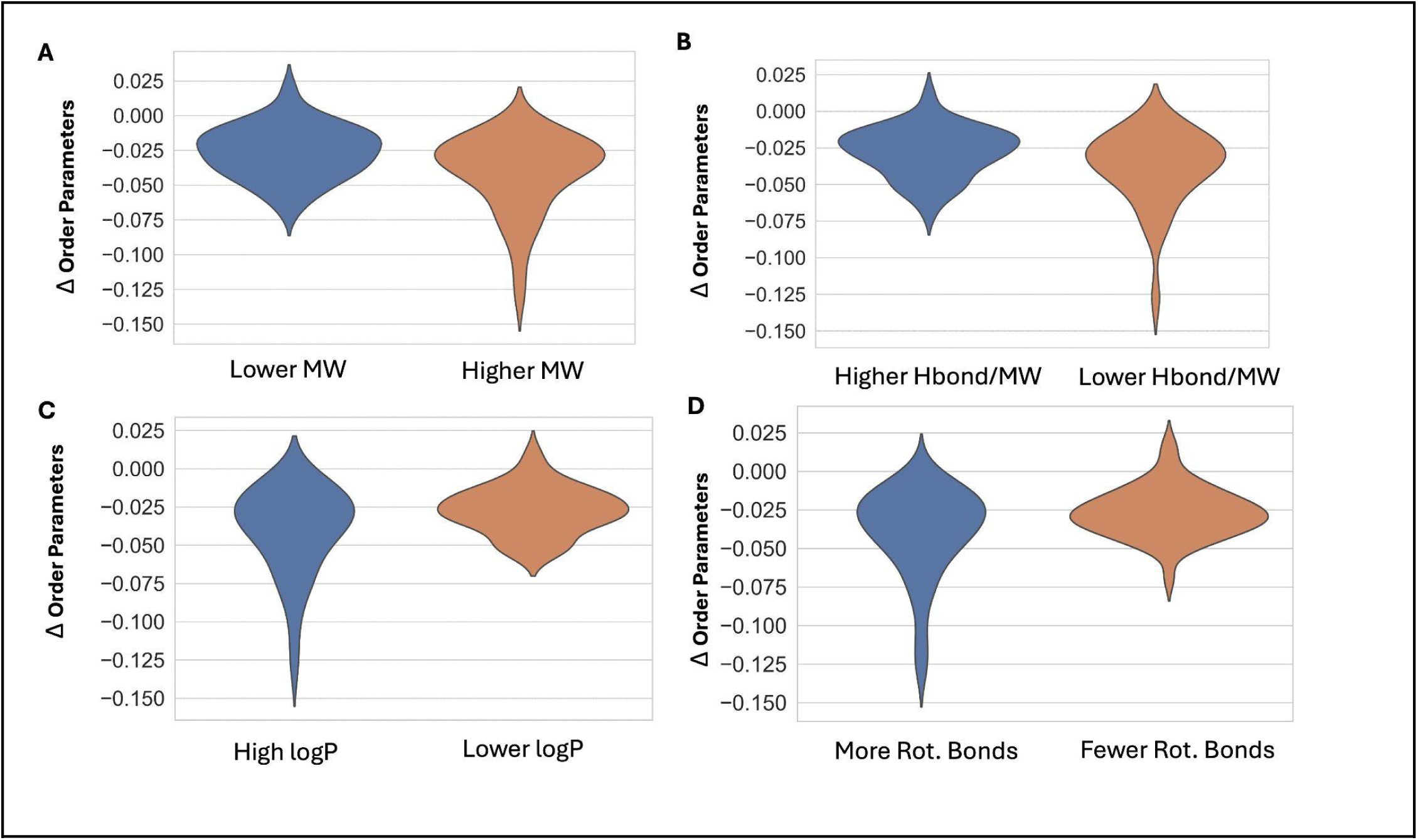
Ligand properties correlation with conformational heterogeneity across the entire protein. **A.** Higher molecular weight is more flexible (Higher molecular weight ≥343.63 Da; ΔOP −0.03, Lower molecular weight ≤ 270.38; ΔOP −0.02; p=0.03; Mann Whitney U test), **B.** More hydrogen bonds/molecular weight is less flexible (More Hbond/MW ≥0.04; ΔOP −0.03, Fewer Hbond/MW≤0.3; ΔOP −0.03; p=3.2e-05), **C.** High logP is more flexible (Higher logP: ≥0.27; ΔOP −0.03, Lower logP: ≤-1.17; ΔOP −0.03; p=0.07) **D.** Higher number of rotatable bonds is more flexible (More rotatable bonds: ≥5; ΔOP −0.03, Fewer rotatable bonds: ≤3; ΔOP −0.02; p=0.07).

**Supplementary Figure 9.**
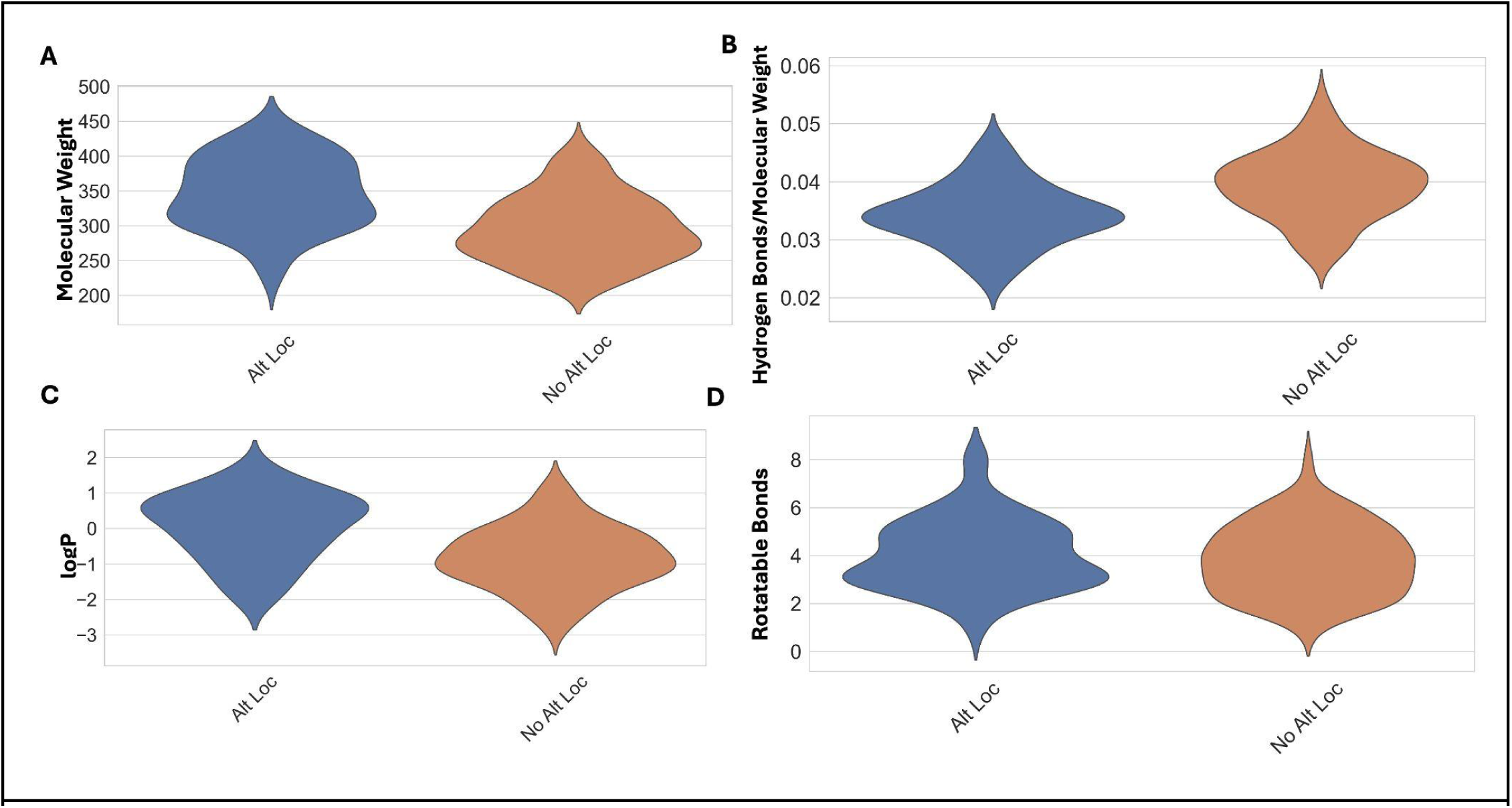
Ligand property association with alternative conformers **A.** Molecular weight (Altloc median=343.9, non-altloc median= 286.91; p=4.39e-08), **B.** Hydrogen Bonds per molecular weight (Altloc median=0.034, non-altloc median= 0.04; p=5.90e-07), **C.** LogP (Altloc median=0.24, non-altloc median=-0.85; p=3.31e-07), **D.** Rotatable bond (Altloc median=4.0, non-altloc median=4.0; p=0.41).

**Supplementary Figure 10.**
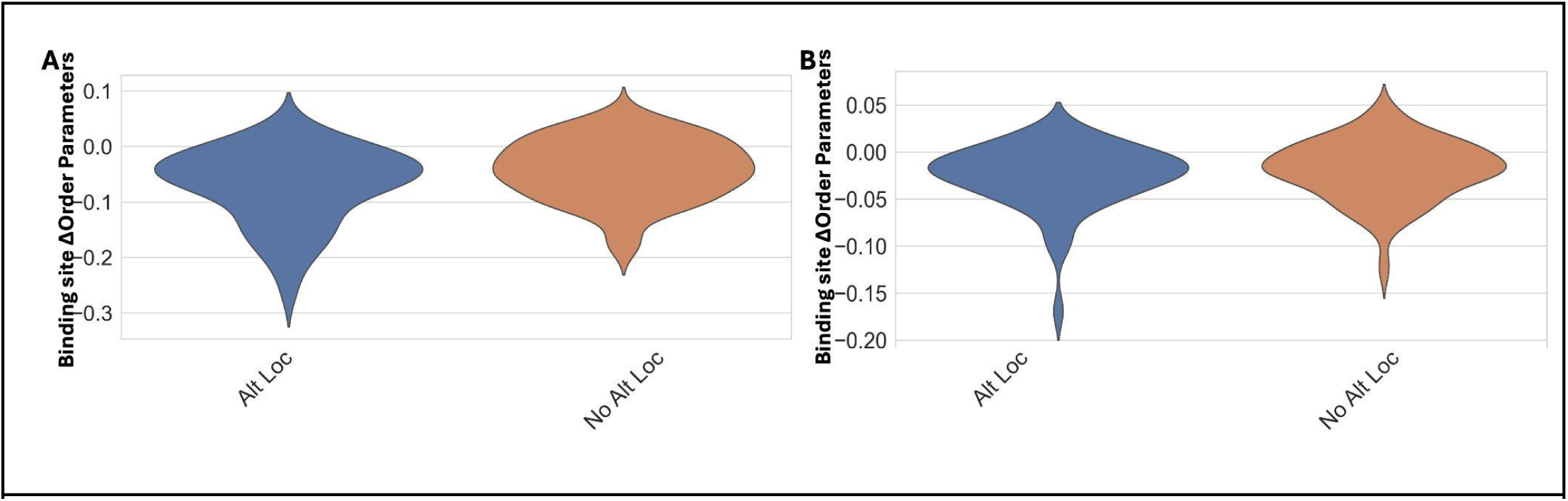
**A.** Binding site order parameter difference (Ligands with modeled alt loc median:-0.059, Ligands without modeled alt loc median: −0.038; p=0.02), **B.** All protein order parameter differences (Ligands with modeled alt loc :-0.019, Ligands without modeled alt loc : −0.018; p=0.33).

**Supplementary Figure 11.**
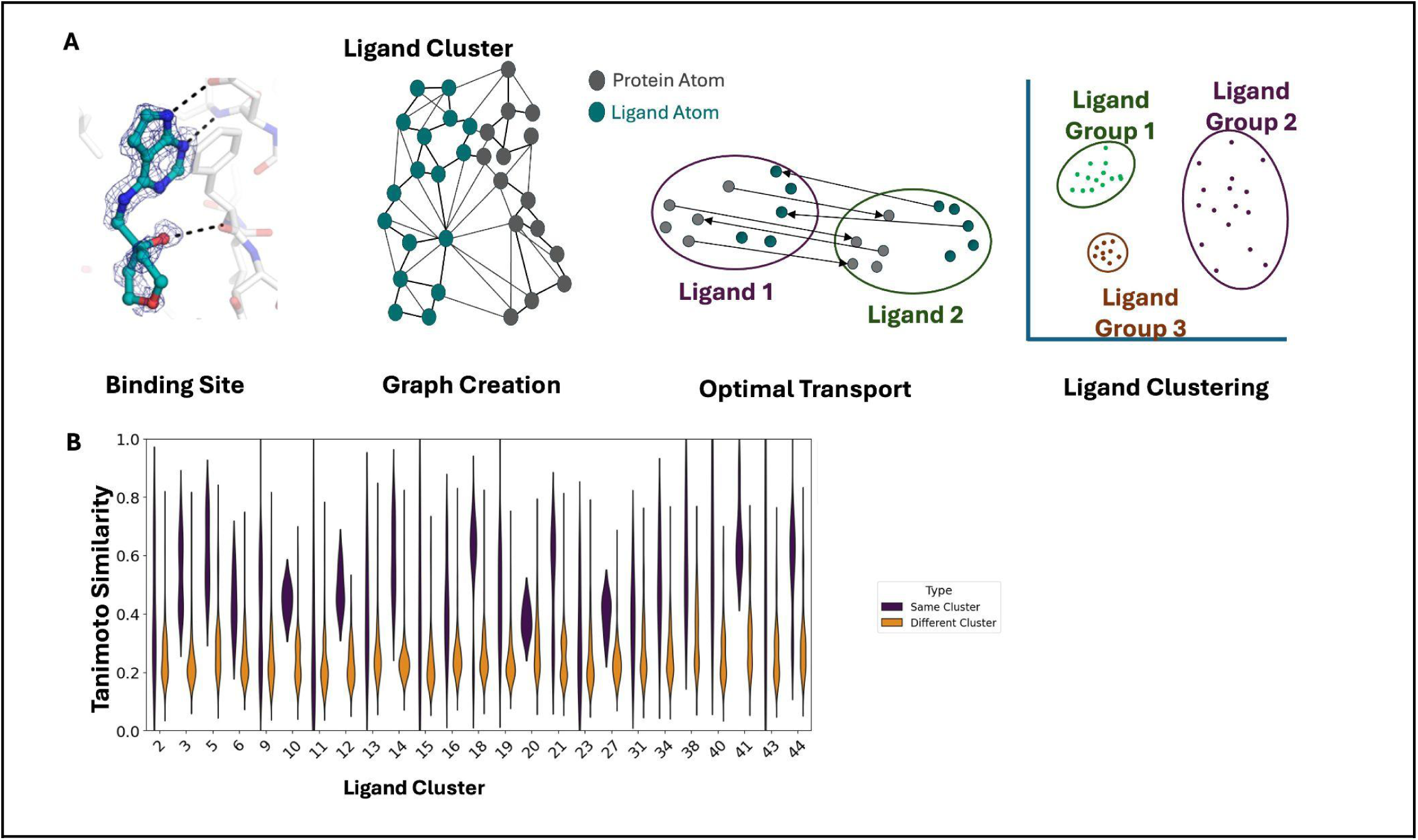
**A**. Protein-ligand interaction optimal transport pipeline. **B.** Tanimoto similarity distribution among ligand clusters.

**Supplementary Figure 12.**
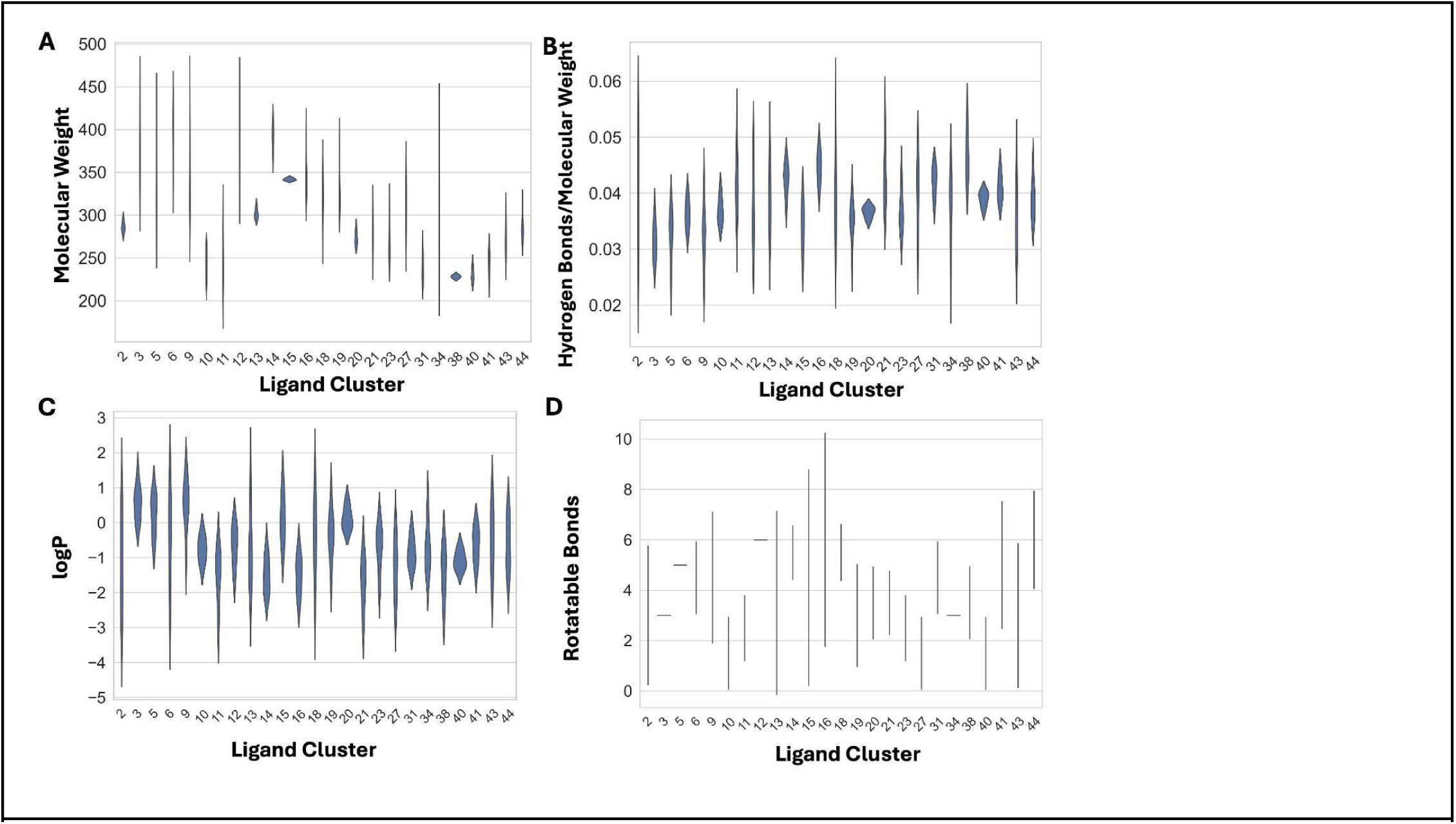
Distribution of ligand properties among ligand clusters.

**Supplementary Figure 13.**
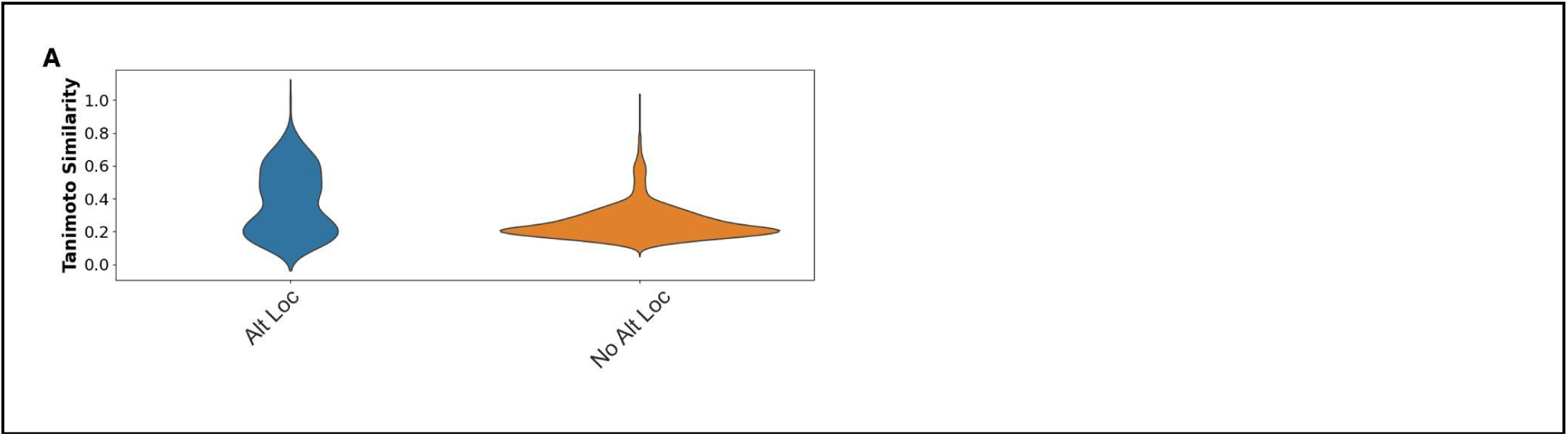
Tanimoto similarity distribution among all ligand with alt locs (blue) vs. ligands without altlocs compared to all ligands with single conformers.

**Supplementary Figure 14.**
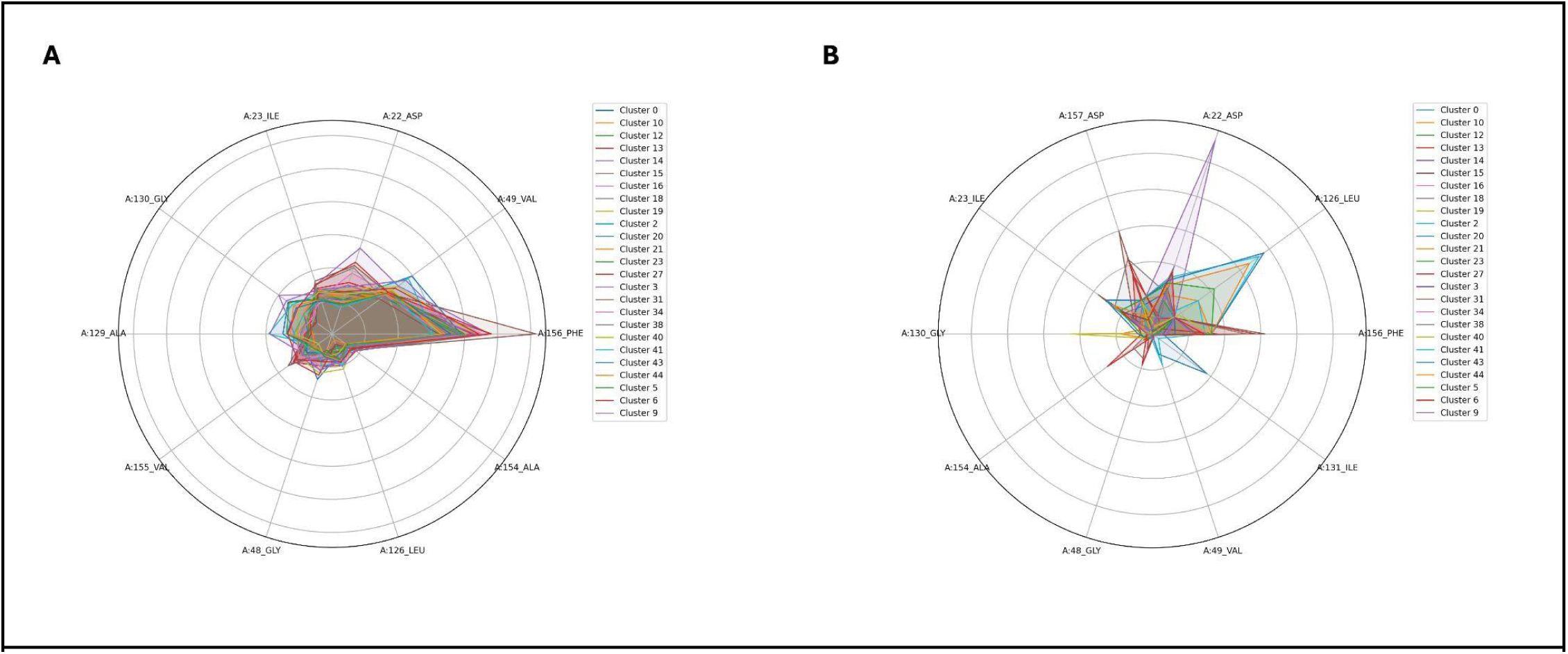
Interaction frequency between binding site residues and ligand groups. **A.** All interactions, **B.** Hydrogen bonds interactions between binding site residues and ligand groups.

**Supplementary Figure 15.**
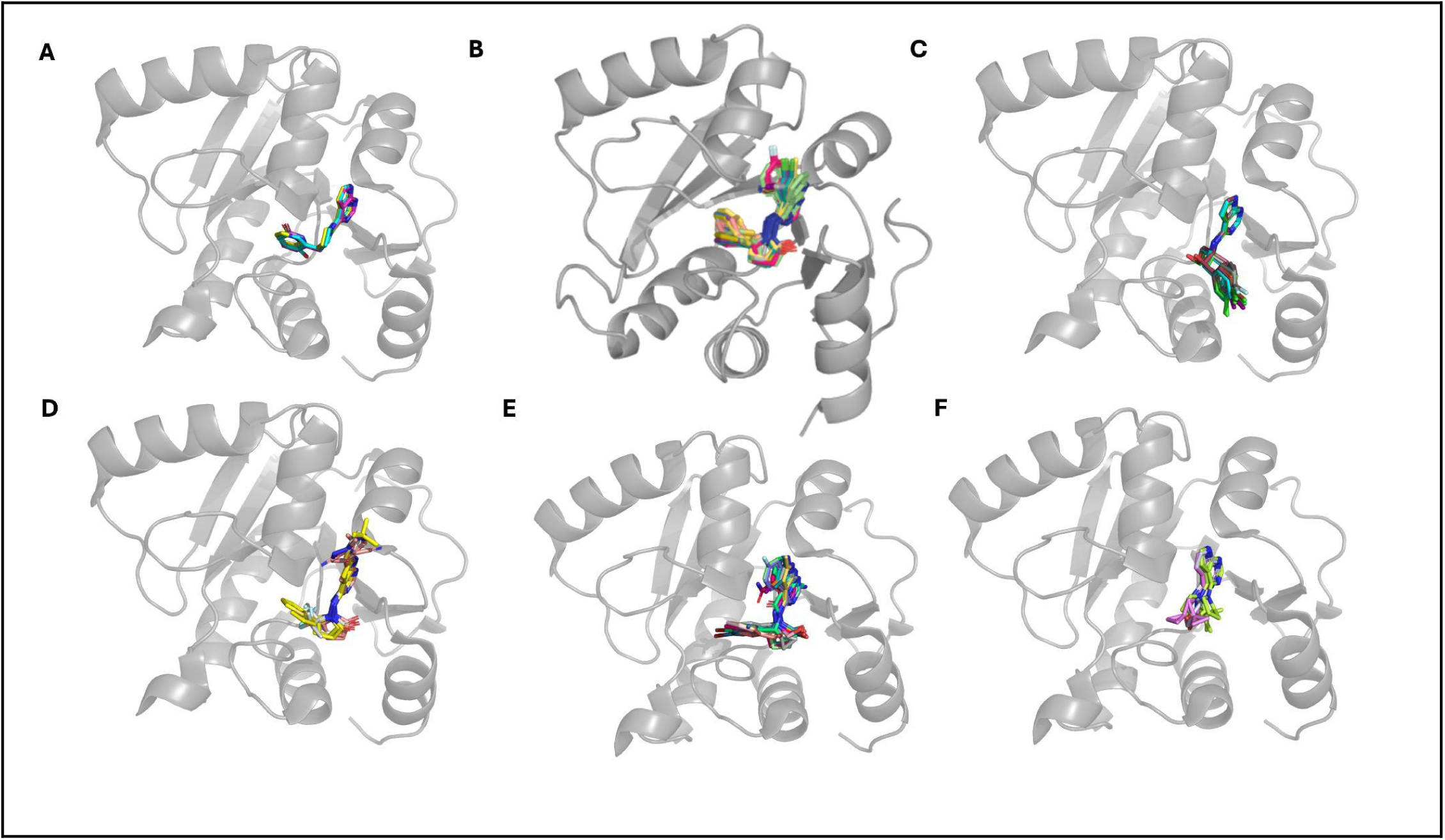
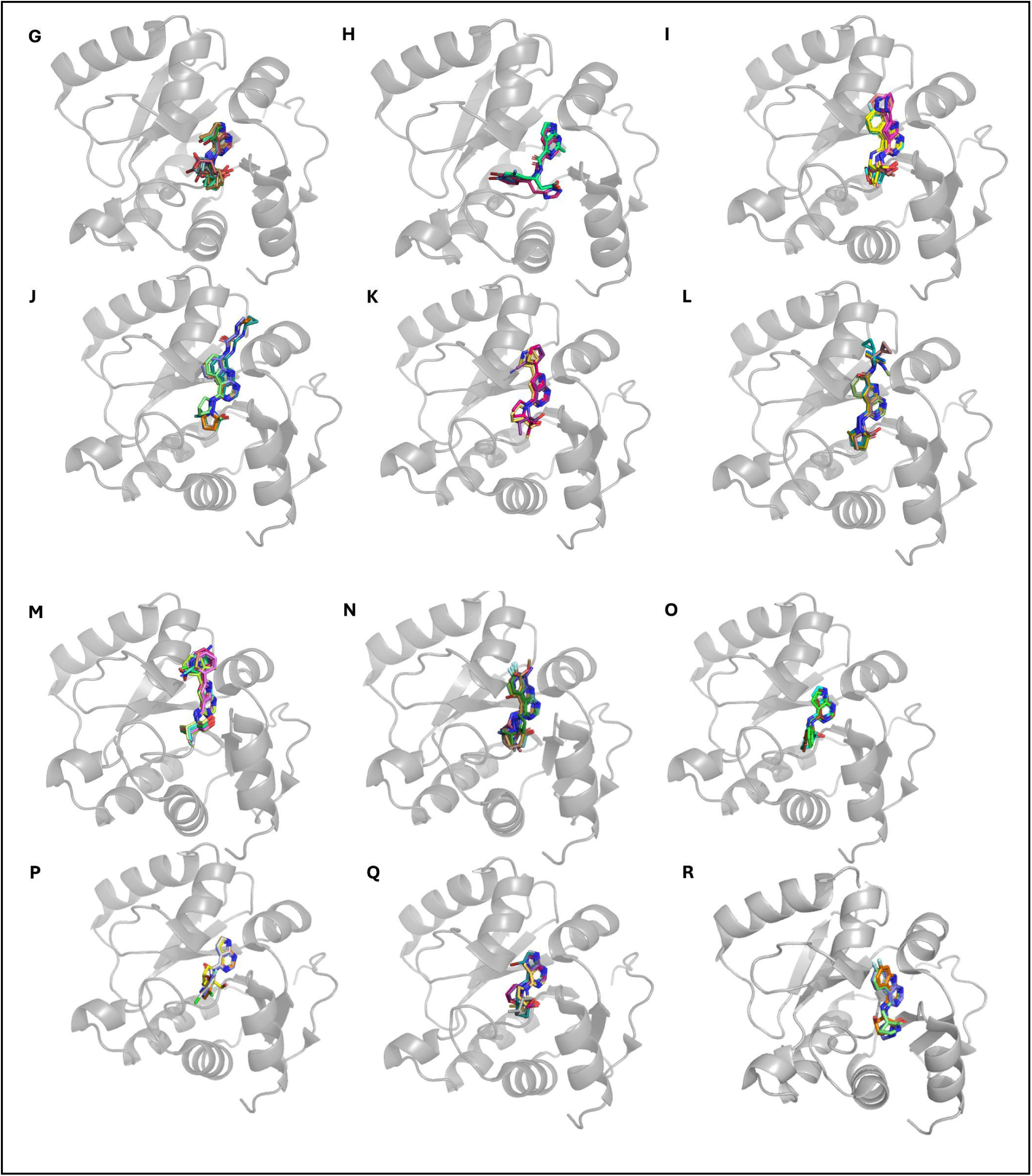
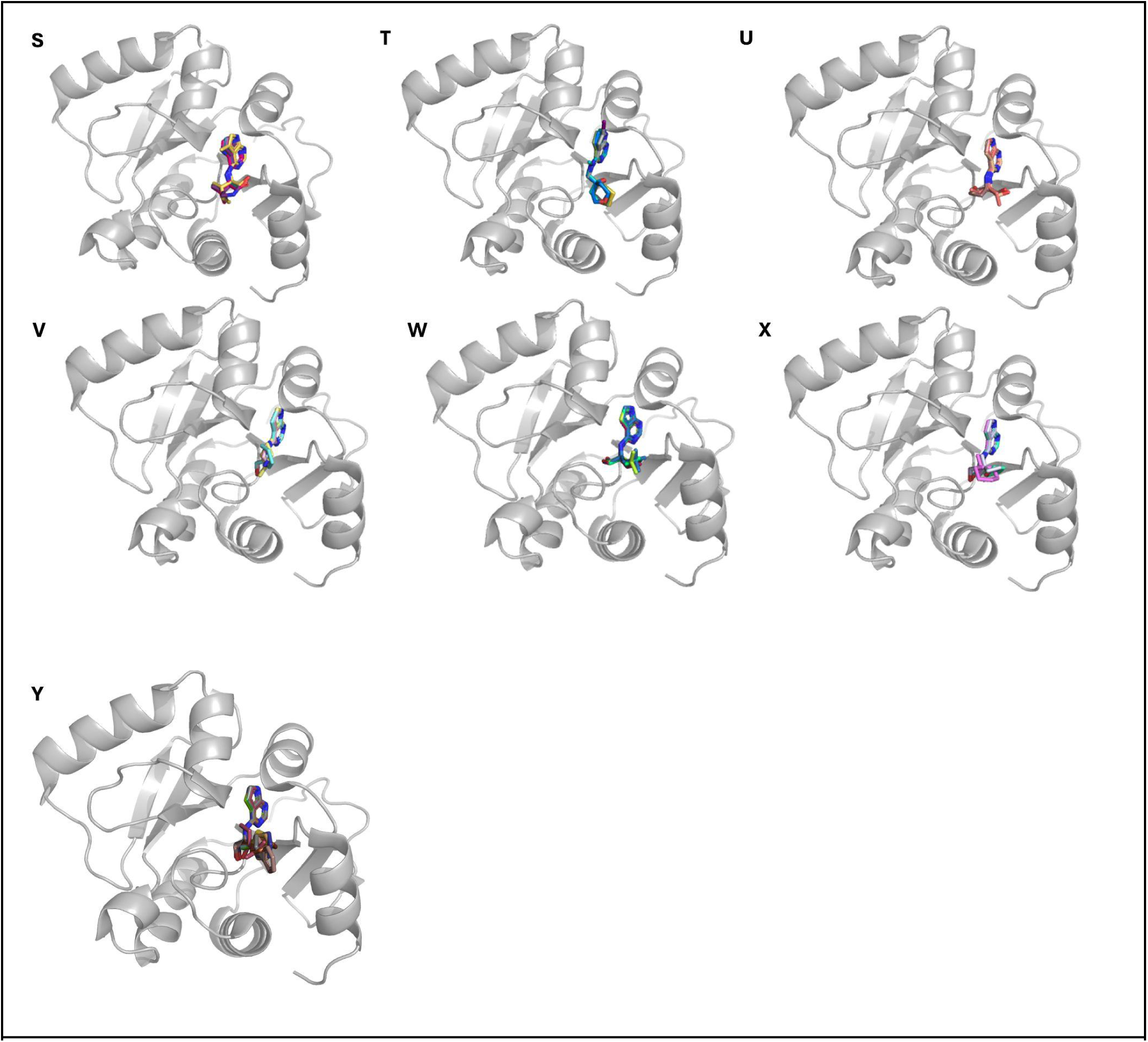
**A.** Ligand Group 2, **B.** Ligand Group 3, **C.** Ligand Group 5, **D.** Ligand Group 6, **E.** Ligand Group 9, **F.** Ligand Group 10, **G.** Ligand Group 11, **H.** Ligand Group 12, **I.** Ligand Group 13, **J.** Ligand Group 14, **K.** Ligand Group 15, **L.** Ligand Group 16, **M.** Ligand Group 18, **N.** Ligand Group 19, **O.** Ligand Group 20, **P.** Ligand Group 21, **Q.** Ligand Group 23, **R.** Ligand Group 27, **S.** Ligand Group 31, **T.** Ligand Group 34, **U.** Ligand Group 38, **V.** Ligand Group 40, **W.** Ligand Group 41, **X.** Ligand Group 43, **Y.** Ligand Group 44

**Supplementary Figure 16.**
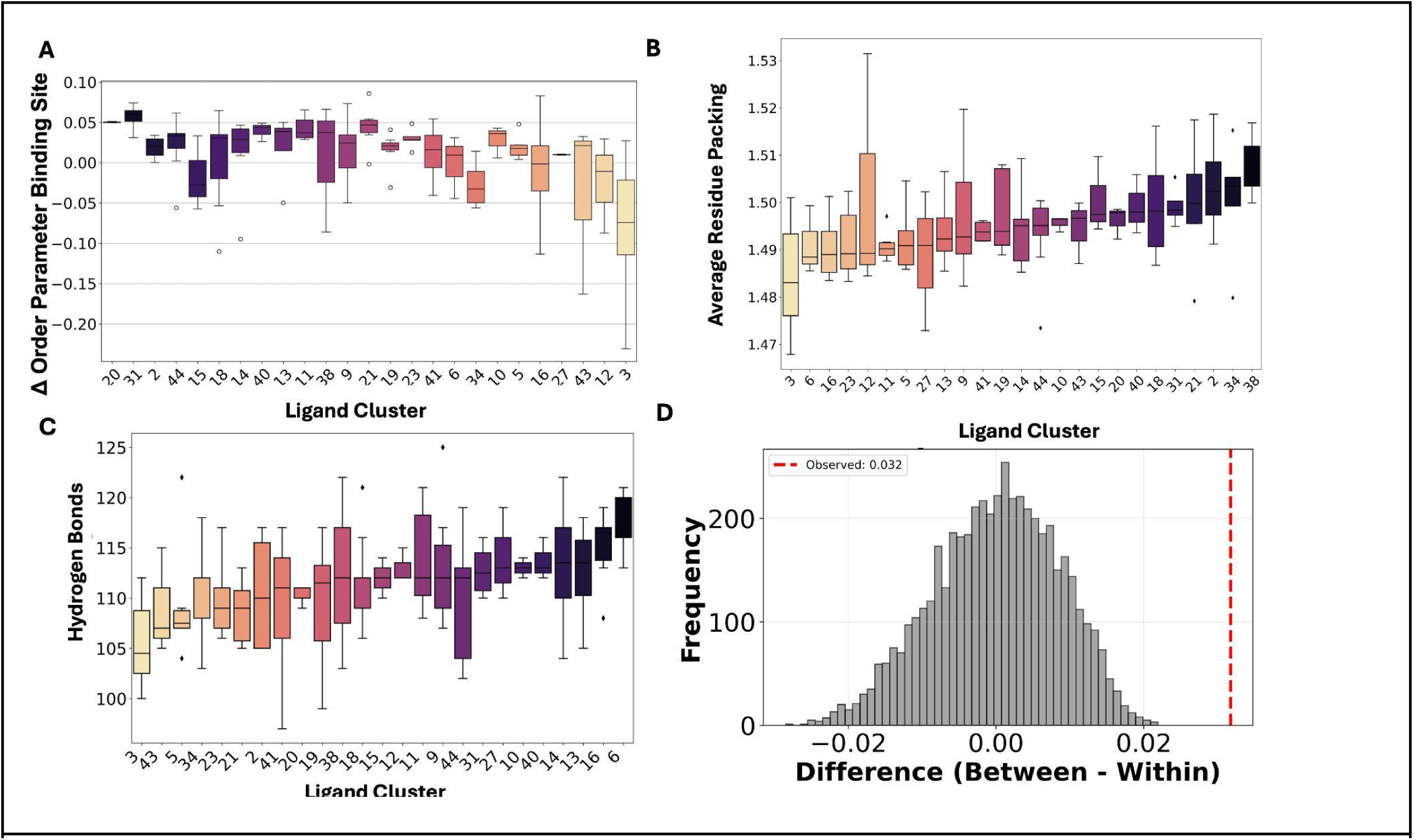
**A.** ΔOrder Parameter in Binding Site per Ligand Group **B.** Packing by ligand group**. C.** Number of hydrogen bonds per ligand group. **D.** Permutation test results from 1-Pearson on order parameter differences.

**Supplementary Figure 17.**
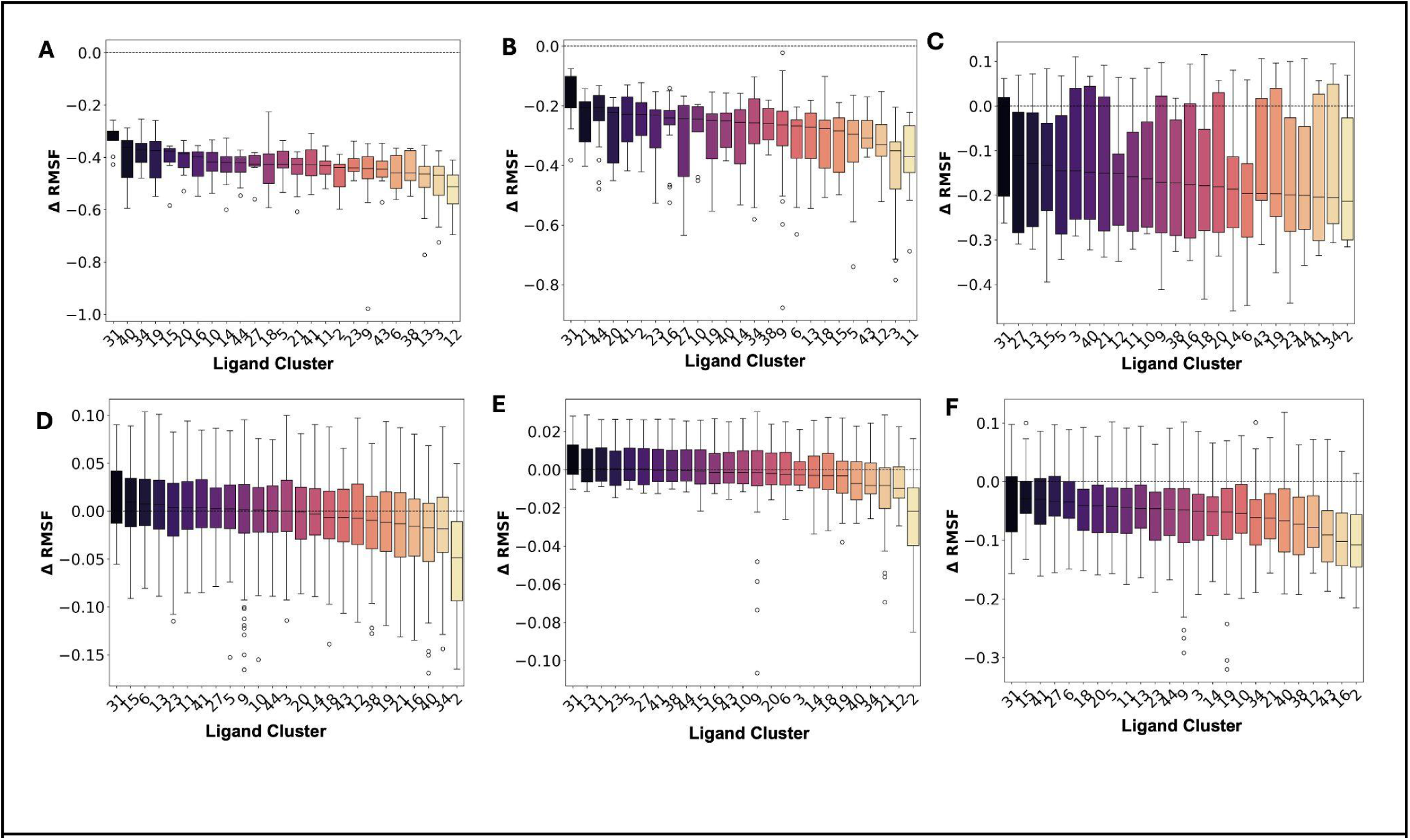
RMSF for each residue cluster separated by ligand group. **A.** Residue Cluster 1 (residue number 102, 103, 104, 105, 106). **B.** Residue cluster 2 (residues 157, 158, 159, 161, 162, 165) **C.** Residue Cluster 3 (residues 29, 30, 31), **D.** Residue cluster 4 (residue numbers 36, 78, 79, 80, 81, 83, 93, 109, 110, 114, 115, 116, 117, 118, 119, 142, 143) **E.** Residue cluster 5 ( 37, 39, 45, 59, 60, 62, 94, 95), **F.** 13, 15, 16, 114, 146, 147, 149, 151, 152.

**Supplementary Figure 18.**
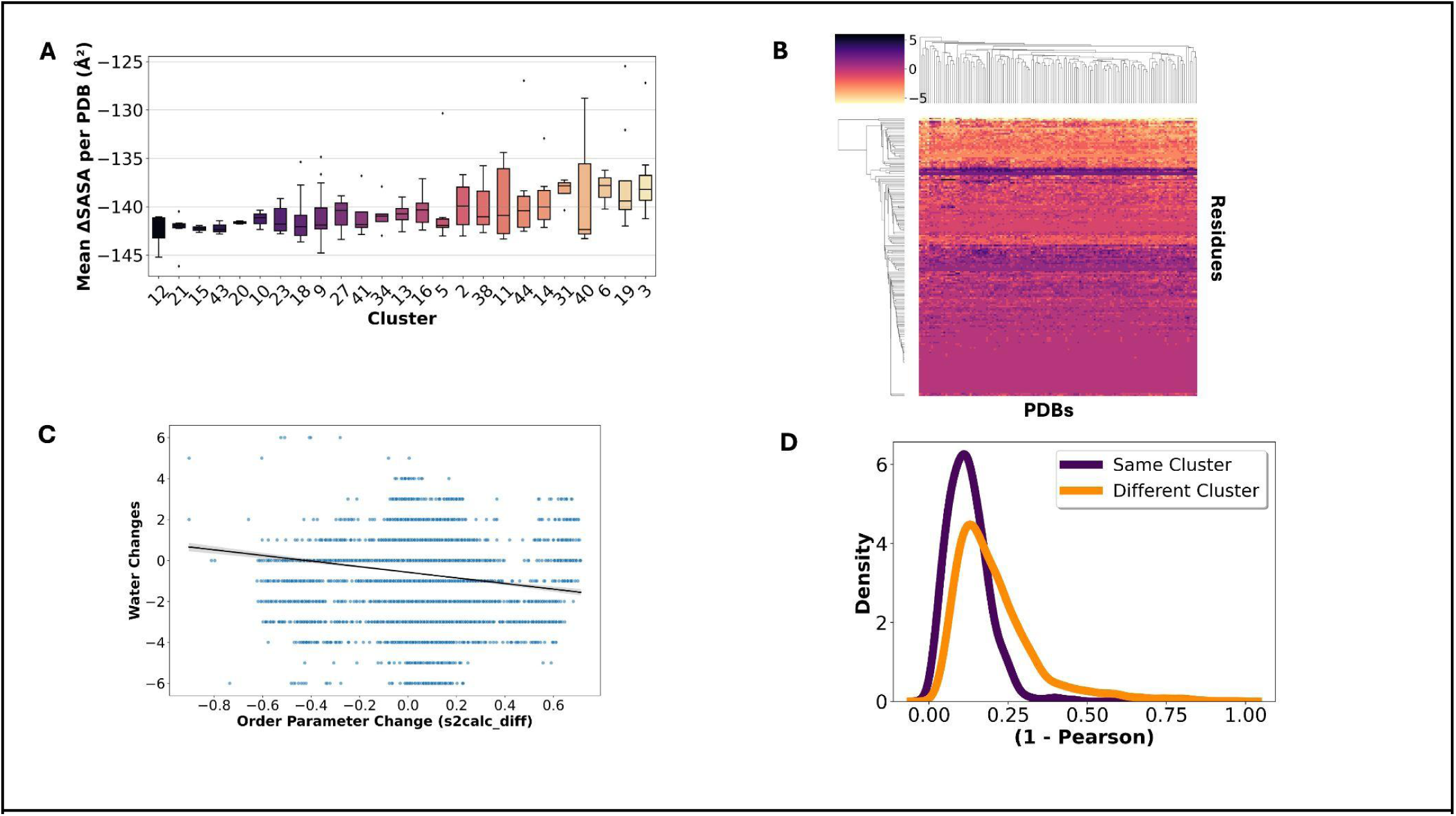
**A.** Change in solvent accessible surface area by ligand group. Negative SASA indicates solvent accessible surface area lost upon ligand binding. **B.** Distribution of water molecules lost or gained across every PDB and every residue. **C.** KDE of water gain or loss exploring only binding site residues defined as any residues with 3 Å of any ligand heavy atom across the dataset (p=1.25e-55; Means: within=0.126, between=0.209).

**Supplementary Figure 19.**
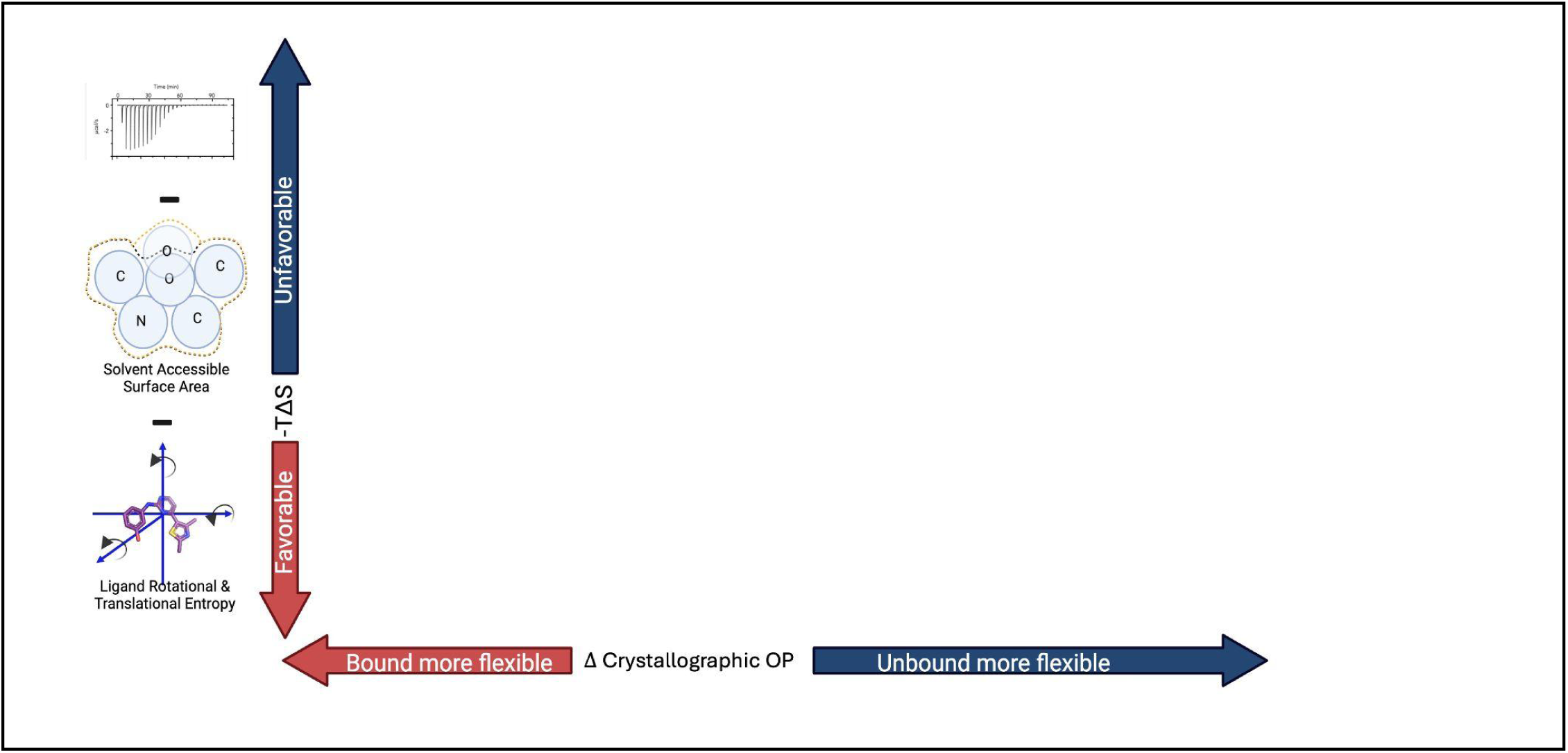
Set up of ITC experiment. On the x-axis is the crystallographic order parameter estimate. On the y-axis is the ITC measured entropy, with solvent and ligand estimated entropy subtracted.

**Supplementary Figure 20.**
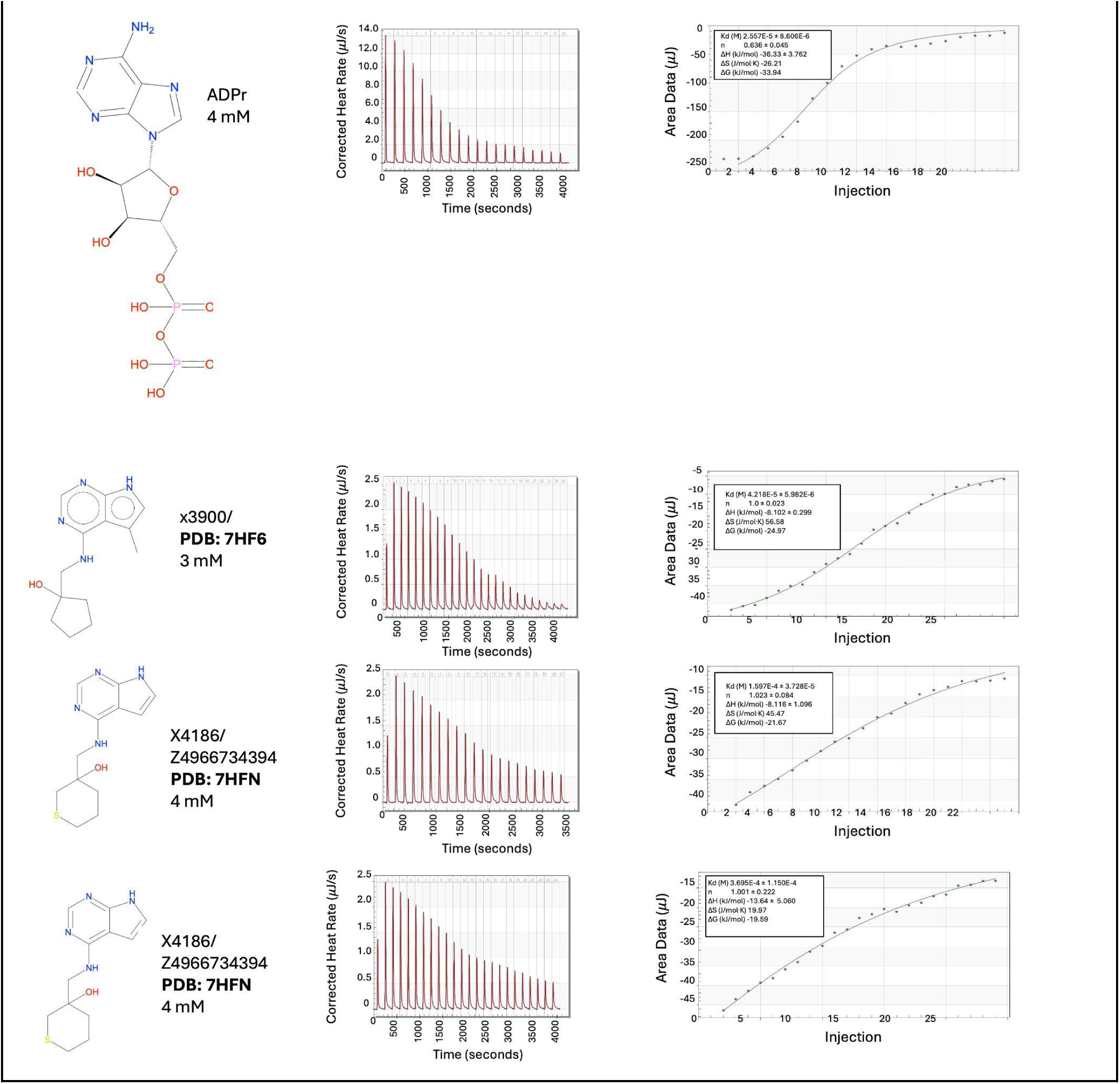
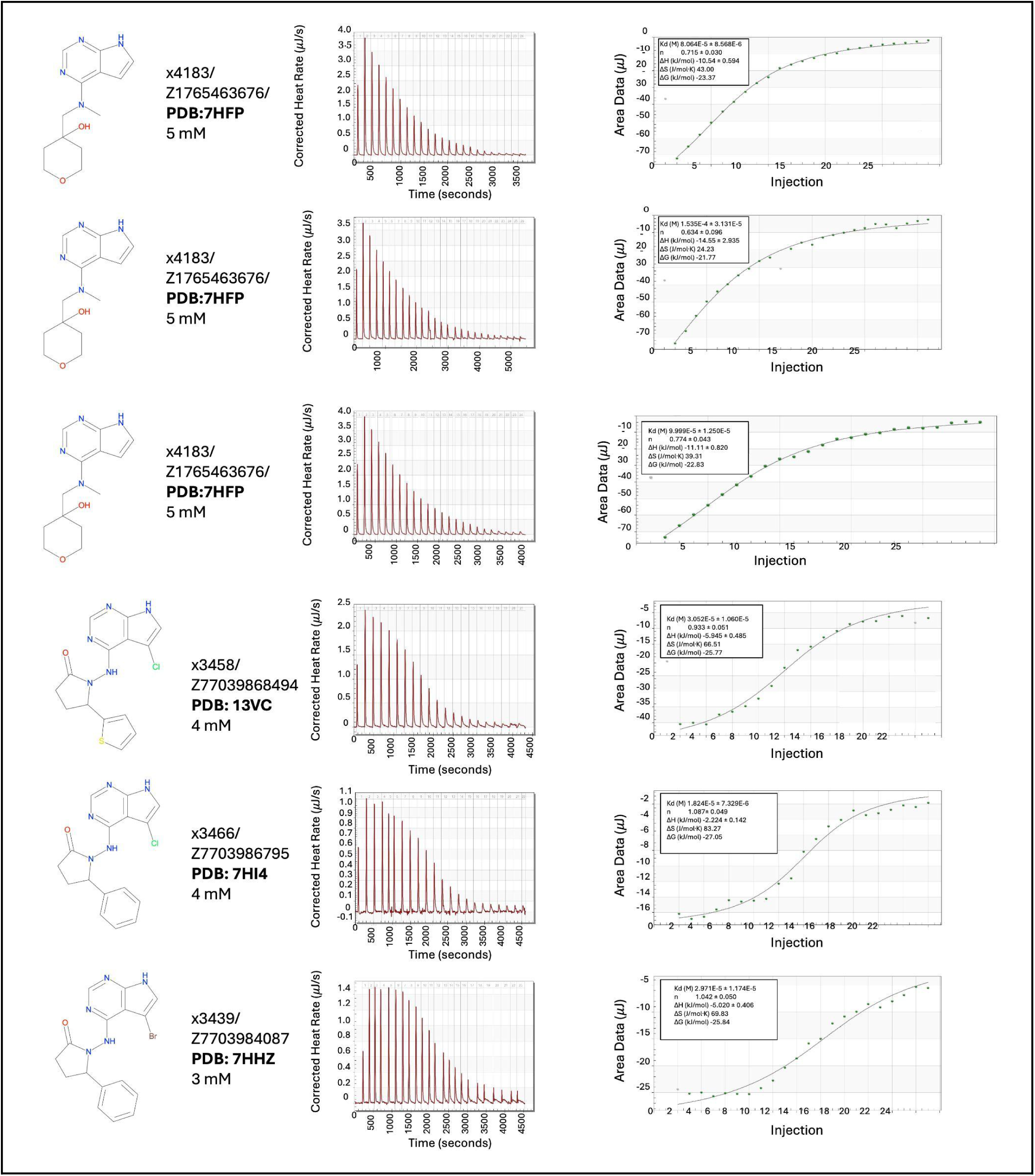
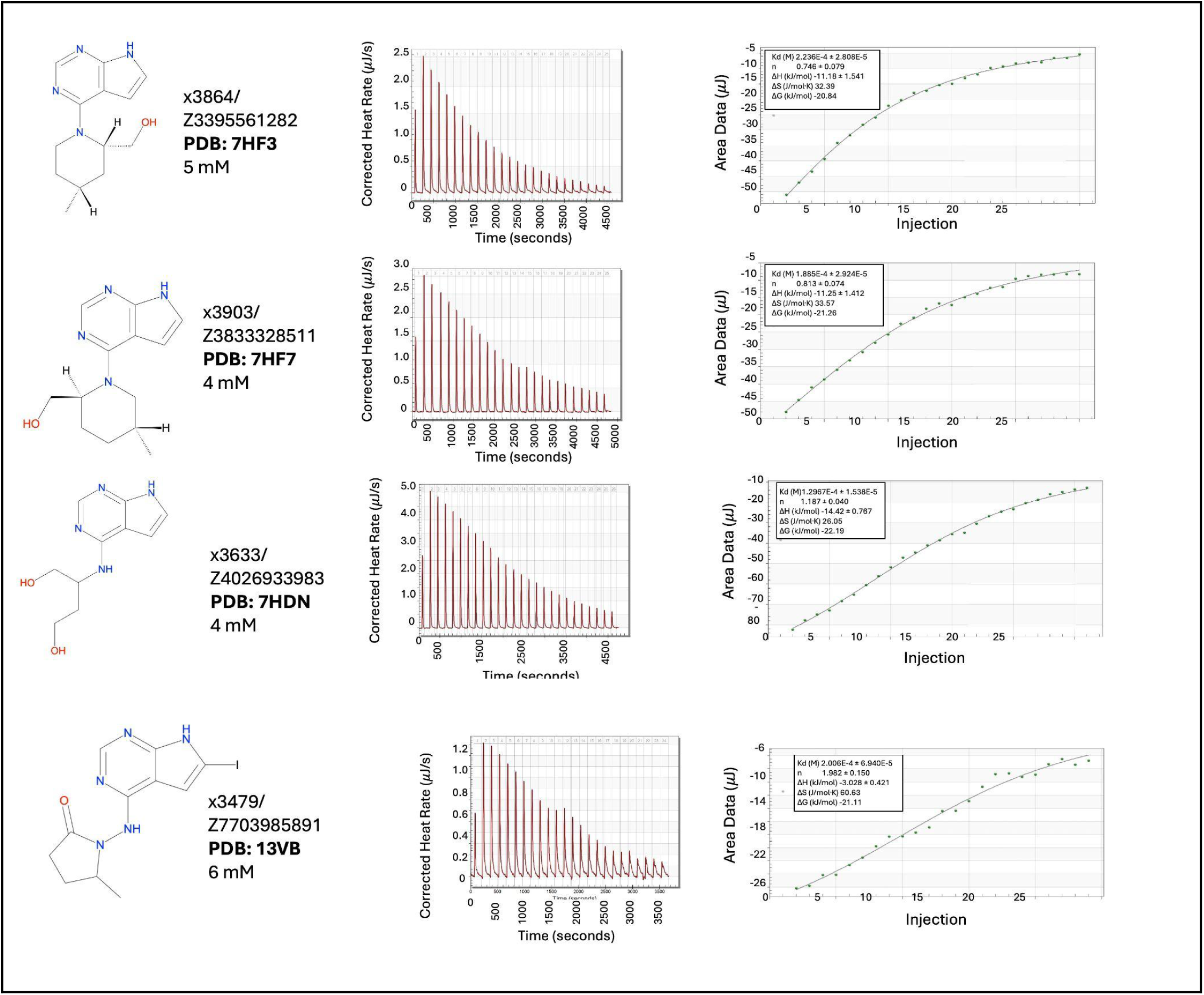
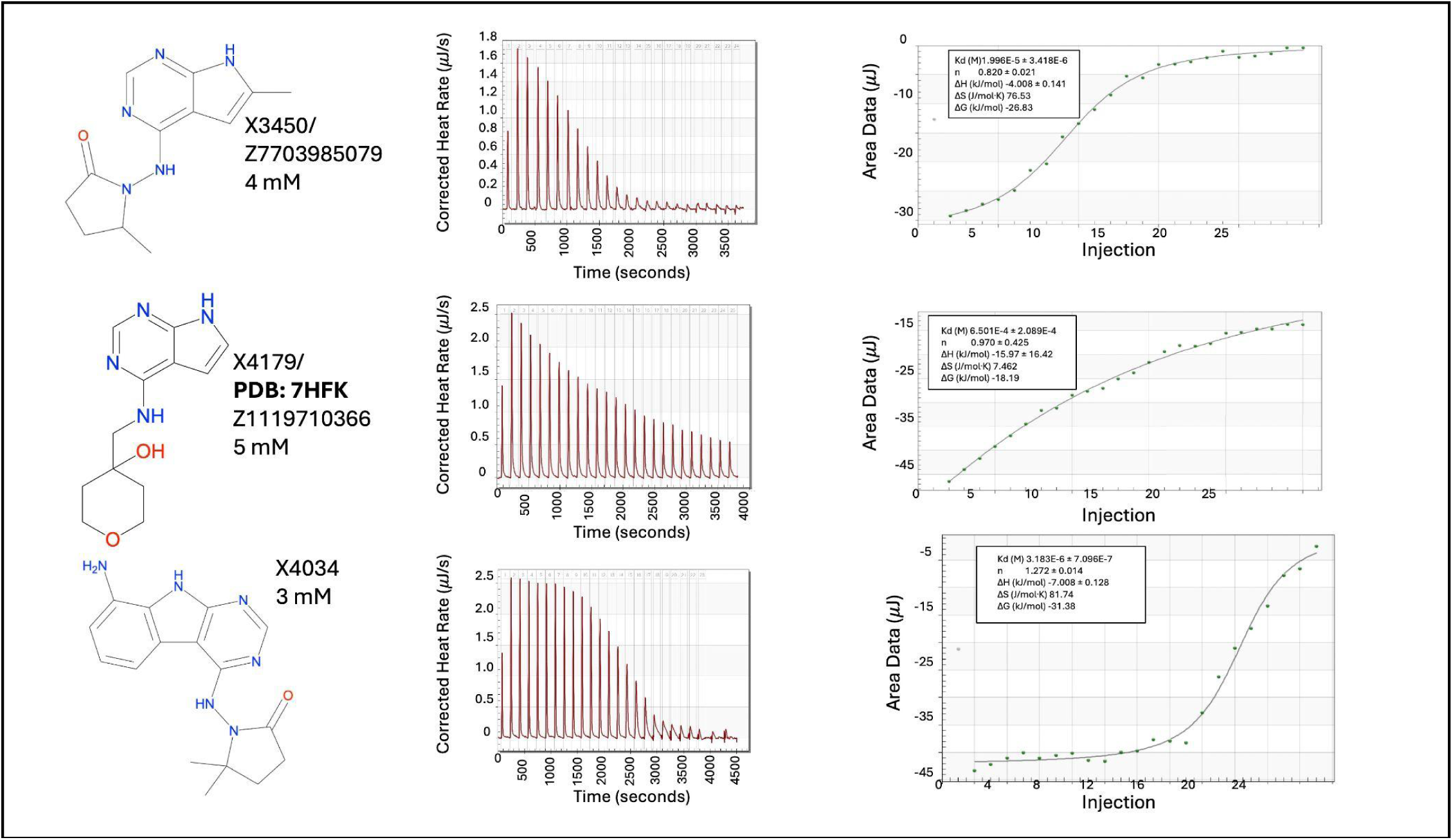
Raw and calculated thermograms for all ligands. If ligands had multiple ITC measurements, we took the average of all runs. Ligands with Z values were ordered from enamine. Others were synthesized at UCSF with descriptions elsewhere.

**Supplementary Figure 21.**
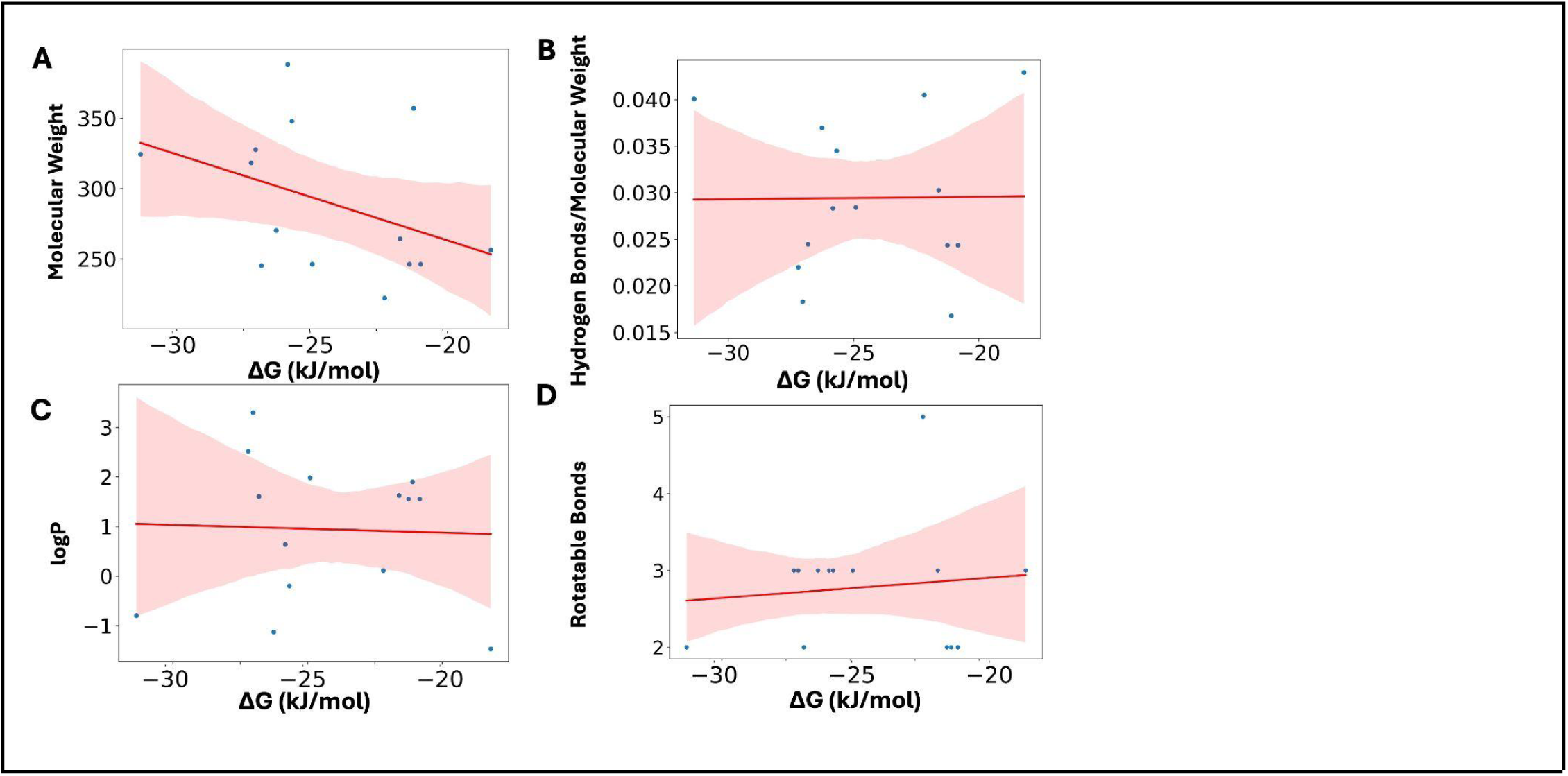
Ligand Properties and Correlation with dG as measured with ITC. A. Molecular Weight (slope:-6.01; R^2^:0.16; p=0.15) B. Hydrogen bonds per molecular weight (slope:0; R^2^= 0.0001; p=0.97). C. LogP (slope:-0.02; R^2^= 0.001; p=0.90) D. Rotatable bonds (slope:0.02; R^2^= 0.01; p=0.71).

**Supplementary Figure 22.**
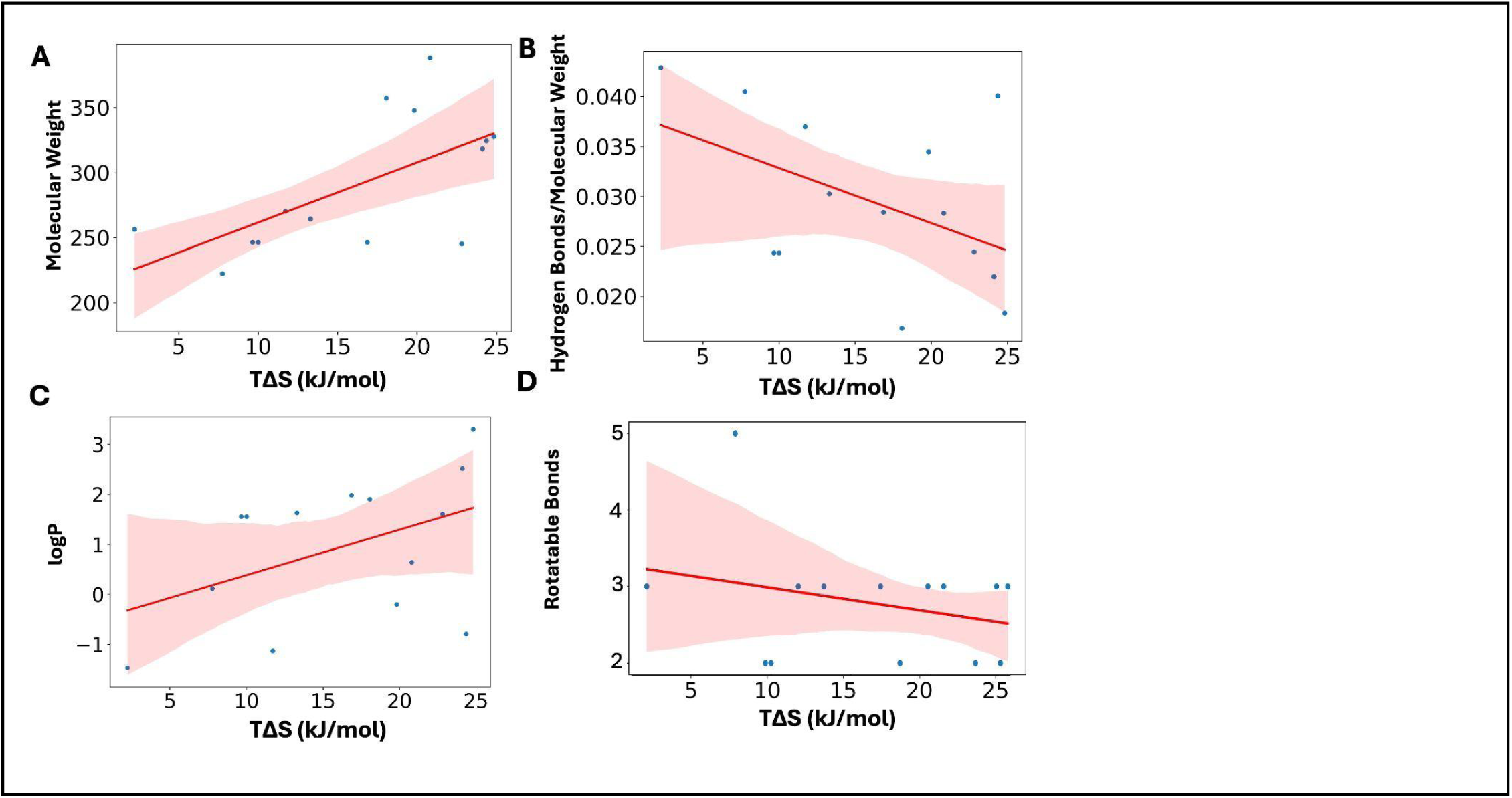
Ligand Properties and Correlation with TΔS. A. Molecular Weight (slope:4.62; R^2^= 0.40; p=0.015) B. Hydrogen bonds per molecular weight (slope:0.038 R^2^= 0.22; p=0.091). C. LogP (slope=0.09; R^2^= 0.20; p=0.11) D. Rotatable bonds (slope=-0.03; R^2^= 0.08; p=0.33).

